# Evaluation of resampling-based inference for topological features of neuroimages

**DOI:** 10.1101/2023.12.12.571377

**Authors:** Simon N. Vandekar, Kaidi Kang, Neil D. Woodward, Anna Huang, Maureen McHugo, Shawn Garbett, Jeremy Stephens, Russell T. Shinohara, Armin Schwartzman, Jeffrey Blume

## Abstract

Many recent studies have demonstrated the inflated type 1 error rate of the original Gaussian random field (GRF) methods for inference of neuroimages and identified resampling (permutation and bootstrapping) methods that have better performance. There has been no evaluation of resampling procedures when using robust (sandwich) statistical images with different topological features (TF) used for neuroimaging inference. Here, we consider estimation of distributions TFs of a statistical image and evaluate resampling procedures that can be used when exchangeability is violated. We compare the methods using realistic simulations and study sex differences in life-span age-related changes in gray matter volume in the Nathan Kline Institute Rockland sample. We find that our proposed wild bootstrap and the commonly used permutation procedure perform well in sample sizes above 50 under realistic simulations with heteroskedasticity. The Rademacher wild bootstrap has fewer assumptions than the permutation and performs similarly in samples of 100 or more, so is valid in a broader range of conditions. We also evaluate the GRF-based pTFCE method and show that it has inflated error rates in samples less than 200. Our `R` package, `pbj`, is available on Github and allows the user to reproducibly implement various resampling-based group level neuroimage analyses.

## 1 Introduction

Mass-univariate inference is a primary tool in localizing regions of the brain associated with a phenotype, disease process, or psychiatric disorder (Yeung, 2018). This approach models the voxel intensity of the 3-dimensional brain image for each subject as a function of covariates, and estimates a set of parameters at each location in the image that quantify the association of the covariates with the imaging outcome (Yeung, 2018; Woo et al., 2014; Friston et al., 1994, 1991). The outcome image can be obtained using MRI to measure blood flow, anatomical features of the brain, or brain activation during a task, and is used to understand average differences between groups or associations with a continous covariate, such as age or disease severity. The parameter estimates and statistics are images that are highly multivariate correlated outcomes with hundreds of thousands of measurements.

Statistical inference in imaging requires estimating the distribution of a topological feature (TF) of the test statistic image, which is, in turn, used to perform inference. Two common examples are inference for local maxima and cluster extent inference (CEI). For local maxima the TF is the height of the statistical image at the local maxima, and for CEI the TF is the size of spatially contiguous clusters above a given statistical threshold. A recently proposed topological features is probabilistic threshold free cluster enhancement (pTFCE), which takes advantage of the spatial correlation of signal in the statistical image, but does not require pre-selecting a statistical threshold (Smith and Nichols, 2009; Spisák et al., 2019).

There are parametric and semiparametric approaches to estimating the distribution of TFs of images. Parametric inference methods for TFs rely on assumptions about the structure of the spatial covariance of the test statistic image in order to approximate the distribution of a TF of the test statistic image (Worsley et al., 1999; Schwartzman and Telschow, 2019). This class of methods includes family-wise error rate (FWER) controlling procedures, such as inference on maxima (Schwartzman and Telschow, 2019) or probabilistic threshold-free cluster enhancement (Smith and Nichols, 2009; Spisák et al., 2019). FWER methods use the distribution of the maximum value of a TF for computing p-values; for example the distribution of the global maximum across the whole image across repeated samplings of the data. We refer to this distribution as the global distribution of the TF. Parametric inference also includes topological false discovery rate (FDR) controlling methods that are included in the SPM software (Chumbley and Friston, 2009; Chumbley et al., 2010; Benjamini and Hochberg, 1995) and first compute unadjusted p-values for local maxima or cluster sizes using their marginal distribution under the global null, and then apply classical FDR controlling procedures to the unadjusted p-values. The marginal distribution is the distribution of the TF across repeated samplings of the data; for example the marginal distribution of a local maximum is the distribution of the value of any local maxima in the statistical image across hypothetical repeated samplings of the data.

In contrast to parametric methods, semiparametric methods use an estimate of the spatial joint distribution of the test statistic image and reduce assumptions about the spatial covariance structure. Permutation and bootstrapping are resampling-based inference procedures that use computational approaches to estimate the null distribution of the test statistic (Winkler et al., 2014, 2016; Vandekar et al., 2019, 2018; Guillaume et al., 2014; Park and Fiecas, 2021; **?**; **?**). While computationally intensive, these resampling-based procedures have received increased attention in neuroimaging as the traditional Gaussian random field theory approximations are increasingly criticized for failing to adequately control error rates at the nominal level (Eklund et al., 2016; Vandekar et al., 2019, 2018; Greve and Fischl, 2018; Silver et al., 2011; Park and Fiecas, 2021).

Most existing tools for semiparametric inference assume exchangeability, which can be violated in neuroimaging data (Vandekar et al., 2019). Robust test statistic images that rely on sandwich covariance estimators to perform inference (Guillaume et al., 2014; Li et al., 2013; Vandekar et al., 2019; White, 1980) yield consistent standard errors when the assumption of exchangeability is violated. It is known that these test statistics are slower to converge than parametric test statistics with univariate and multivariate outcomes, but their performance has not been rigorously evaluated in finite samples in neuroimaging data. Previous papers evaluating the effect of heteroskedasticity on resampling methods for robust and parametric test statistics were restricted to unidimensional parameters (T-statistics) (Vandekar et al., 2019) and did not evaluate the efficacy of resampling methods for TFs use in neuroimaging (Guillaume et al., 2014).

In this paper, we evaluate several resampling methods for inference of multidimensional (F-statistics) robust and parametric statistical images in neuroimaging data for marginal and global distributions for several different TF of the statistical image using realistic simulations. We use extensive bootstrap-based simulations to show that the gold-standard permutation procedure with parametric test statistics fails to control the FWER under heteroskedasticity for many commonly used TFs. Simply using robust test statistics with the permutation resampling procedure resolves the issues with this standard permutation approach. The semiparametric bootstrap has close to nominal error rates for parametric and robust test statistics. In addition, we compare these resampling methods with the recently proposed parametric pTFCE approach (Spisák et al., 2019). Finally, we use the methods to study sex differences in the life-span curves of gray matter volume measured with voxel-based morphometry (VBM) in the Nathan-Kline Rockland sample (Nooner et al., 2012). Our findings suggest that using robust test statistics as the standard in neuroimaging may produce more reproducible results.

## 2 Model setup

We consider group-level analysis where there is a single image for each subject: let *Y*_*i*_(*v*) denote the outcome image for subject *i* = 1, …, *n*, which measures a biological or physiological value, such as brain activation to a task, blood flow, resting connectivity with a target brain region, or structural features in gray or white matter. All images are indexed by the voxel location, *v ∈* 𝕍. We assume that (Vandekar et al., 2019)

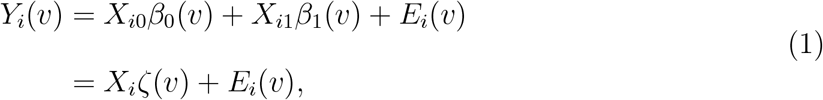

where 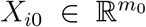 is a row vector of nuisance covariates including the intercept, 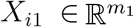 is a row vector of variables of interest, *m* = *m*_0_ + *m*_1_, *X*_*i*_ = [*X*_*i*0_, *X*_*i*1_], *β*_0_(*v*), and *β*1(*v*) are parameter image vectors that take values in 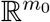 and 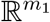, respectively; *β*(*v*) = [*β*_0_(*v*)^*T*^, *β*_1_(*v*)^*T*^]^*T*^, and *E*_*i*_(*v*) is an error term with 𝔼{*E*_*i*_(*v*)} = 0 and spatial covariance function ∑_*i*_(*v, w*) = Cov {*E*_*i*_(*v*), *E*_*i*_(*w*)} < ∞ for any two voxels *v, w*. This covariance depends upon the subscript *i* intentionally to account for differences in the spatial covariance of any two given regions between subjects. Here, all capital letters denote random variables and we require that Cov(*X*_*i*_) is positive definite. Lastly, let 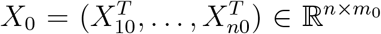, 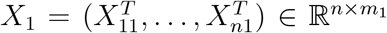, *X* = [*X*_0_, *X*_1_], *Y* (*v*) = [*Y*_1_(*v*), …, *Y*_*n*_(*v*)]^*T*^. In our previous paper we considered weighted least squares regression; to simplify notation here, we consider unweighted least squares estimation, but the weighted least squares of an outcome imaging data 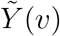 onto covariates 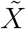 with weights *W* (*v*) is equivalent to least squares with 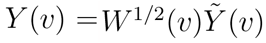 onto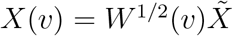. It is also possible that *X* depends on location *v*, but resampling in this case becomes very computationally intensive, so is not discussed here.

Throughout this paper, we discuss estimation of the joint distribution of a test statistic image, *T*_*n*_(*v*), in finite samples, where *T*_*n*_(*v*) is the test statistic image of the null hypothesis

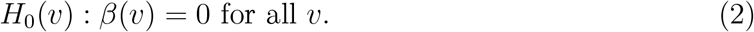

The alternative is, most generally *H*_1_ : *β*(*v*)*≠* 0 for at least one voxel *v*. The hypothesis (2) is tested indirectly by computing *p*-values for TFs of *T*_*n*_(*v*), such as the maximum statistic across the image or the maximum cluster size.

## 3 Methods

### 3.1 Resampling procedures

We describe each of the resampling methods in terms a vector-valued image of “standardized parameters”, *Z*_*n*_(*v*) that are a function of the parameter estimators and their variance estimators,

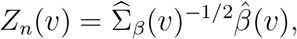

where 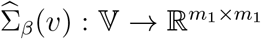 is the estimated covariance of the parameter estimates for each location *v*. The sum of squares of the standardized parameters, *Z*_*n*_(*v*) is proportional to the test statistic image

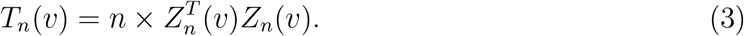

TFs of the test statistic image, *T*_*n*_(*v*) are used for testing the hypothesis (2).

Details to derive *Z*_*n*_(*v*) are given in the Supplement. Briefly, let *A* and *B*(*v, w*) be the “bread” and “meat” components of the sandwich covariance matrix for the estimator image 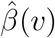 (Vandekar et al., 2019; Boos and Stefanski, 2013; Huber, 1964). *A* only depends on the design matrix *X*, so is not a function of voxel locations, *v* and *w*. We denote their estimators with a hat, such as 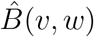. These matrices are derived in the context of M-estimation, where the matrix *A* is the limit of the second derivative of the least squares estimating equation (given in the supplement) and *B* is the limit of the expected value of the outer product of the first derivative of the least squares estimating equation. Please see our previous paper (Vandekar et al., 2019) and the supplement for a more detailed description. When *v* = *w* we write *B*(*v, v*) = *B*(*v*) to simplify notation. 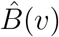 depends on *v* only through *Y* (*v*), so we will write *B*(*v*) = *B*{*Y* (*v*)} at times, to express it as a function of *Y* (*v*). Using this notation, the standardized parameter images are

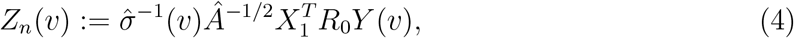

for the parametric test statistic and

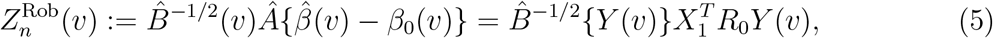

for the robust test statistic. Note, *B*^*−½*^{*Y* (*v*)} is an *m*1 *× m*1 matrix valued function of *v*, so we can easily compute this inverse square root matrix at each location *v*. *B*{*Y* (*v*)} is the covariance of the parameter 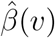 and does not describe the spatial dependence of 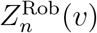, which is estimated in Sections 3.1.1 and 3.1.2 via resampling. The distinction between (4) and (5) is synonymous to the difference between univariate test statistics computed using likelihood-based covariance estimators and heteroskedasticity consistent estimators (see e.g. Boos and Stefanski, 2013, Ch. 7).

The notation we introduce here allows us to easily describe how each of the resampling procedures computes the standardized parameters, *Z*_*n*_(*v*), using the outcome images, *Y* (*v*), which gives us the joint distribution of the test statistic image *T*_*n*_(*v*) through (3). Thus, our goal to estimate the distribution of TF of the test statistic image can be achieved by reproducing the joint distribution of *Z*_*n*_(*v*).

Many resampling procedures to estimate the joint distribution of *Z*_*n*_(*v*) can be expressed in terms of (4) or (5). Throughout this section we let 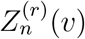 denote an arbitrary resampled version of the parametric or robust estimator.

#### 3.1.1 Permutation procedure

The Freedman-Lane permutation procedure is the most commonly used permutation procedure in neuroimaging (Winkler et al., 2014, 2015, 2016; Nichols, 2012). It assumes exchangeability conditional on the mean under the null (2),

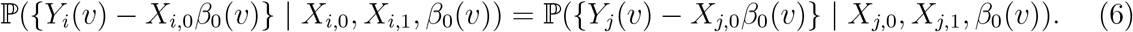

Since *β*_0_(*v*) is unknown, the method conditions on an estimate of *β*_0_(*v*) and draws a sample from the joint distribution of *Z*_*n*_(*v*) using a random permutation matrix *P* (Winkler et al., 2014),

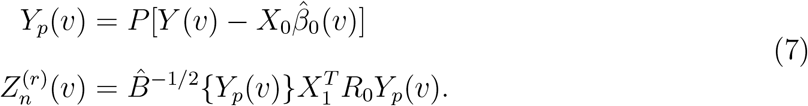

Note that, because the permutation breaks any possible dependence between the variance of *Y* (*v*) and *X*_1_, it does not accommodate heteroskedasticity; this approach may reject if the null (2) is true, but (6) is not.

#### 3.1.2 Wild bootstrap methods

We consider wild bootstrap methods to sample the outcome images, *Y* (*v*), and compute the test statistics (Wu, 1986; Efron and Tibshirani, 1994). The wild bootstrap preserves the association between the residuals and the covariates so that it reproduces unequal variance if it exists in the population. The procedure computes the resampled test statistic,

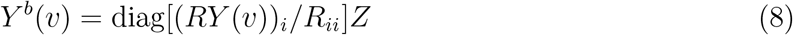

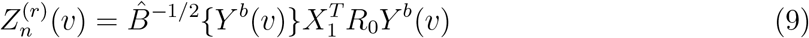

where *Z ∈* ℝ^*n*^ is a random variable generated computationally. We consider two wild bootstrap methods based on methods that were effective in similar settings (Bowring et al., 2019; Sommerfeld et al., 2018; Djogbenou et al., 2019; Guillaume et al., 2014): the first samples the elements of *Z* as independent normal random variables and the second samples the elements of *Z* as independent Rademacher random variables, which take the values -1 and 1 with equal probability. This is equivalent to sign flipping the residuals, so assumes symmetric errors, but does not assume exchangeability because the residuals are not permuted (Winkler et al., 2014). Dividing by the diagonal of the residual forming matrix, *R*_*ii*_, is a jackknife approximation that adjusts for the bias of the squared residual (Long and Ervin, 2000; MacKinnon and White, 1985). Our previous approach (Vandekar, 2019) conditioned on the covariance matrix:

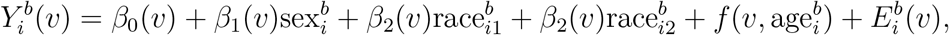

where *Y* ^*b*^(*v*) is as defined in (8). In simulations, we have found this procedure does not accurately reproduce the distribution of the test statistics in small samples. Bowring et al. (2019) also found that conditioning on the covariance estimator does not work well. Instead, our approach here is similar to a wild t-bootstrap (Bowring et al., 2019; Telschow and Schwartzman, 2019), and simplifies to the wild t-bootstrap using the parametric test statistic (4). For all procedures, we apply a T to Z quantile transformation to *Z*_*n*_(*v*), which we have found improves the convergence rate of the test statistics *T*_*n*_(*v*) to their asymptotic distribution (Vandekar et al., 2019).

### 3.2 Topological features of the test statistic image

We are interested in estimating the distribution of TF of the test statistic image in order to perform one of several potential inference procedures. We consider three types of TF: local maxima, cluster extent, and cluster mass Spisák et al. (2019). We use a nonbinary dilate algorithm to define local maxima: The pixel value at a given location in the dilated image is assigned the maximum value within a 7 voxel (14mm) window around the center voxel. We define local maxima as any pixel in the dilated image that equals the value in the original image. We used this algorithm because it does not require computation of an image dertivative to determine maxima.

CEI and cluster mass inference (CMI) depend on the cluster forming threshold, which we choose to satisfy an uncorrected voxel-wise threshold of *α* ∈ {0.01, 0.001}. These inference procedures first threshold the image using the 1 *− α* quantile of the reference distribution, 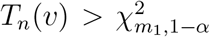, where 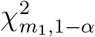 is the 1 *− α* quantile of the chi-squared distribution with *m*1 degrees of freedom. Then, we determine spatially contiguous clusters using a connected components algorithm. The CEI statistic for each contiguous cluster is the number of voxels included in the cluster, and the CMI statistic is the sum of the statistical image values in each cluster.

The marginal distribution is used to compute uncorrected p-values for each of the TF. We compute the marginal distribution using all of the local maxima or clusters from each bootstrap. For *P* bootstrap or permuted samples, the marginal distribution of the cluster size is estimated by

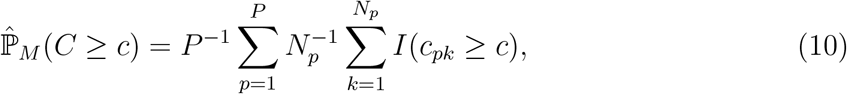

where *C* denotes the random variable that has the marginal distribution of the local maxima or cluster extent/mass, *N*_*p*_ is the number of local maxima or clusters in bootstrap *p, c*_*pk*_ is the *k*th maxima or cluster value in bootstrap/permutation *p*, and *I* is the indicator function. We then apply the Benjamini-Hochberg adjustment to the vector of p-values for the observed clusters (Benjamini and Hochberg, 1995).

The global distribution is used to compute FWER-adjusted p-values. The global distribution is the distribution of the maximum voxel value, maximum cluster size, and maximum cluster mass, for each of the TF.

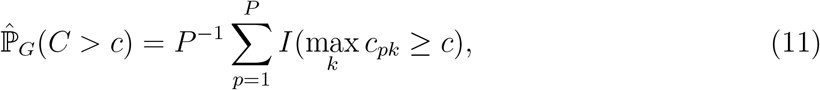

where objects are as defined for (10). These TF were chosen because they apply to most commonly used inference procedures and can be used for FWER and FDR control of voxelwise inference, CEI, and CMI, and also for constructing spatial confidence sets.

pTFCE is a topology-based approach that is applied to the statistical image to compute a new statistical image that aggregates spatially localized statistical values across a range of CFTs to improve sensitivity to diffuse and localized regions of signal (Spisák et al., 2019). The standard approach for inference uses Gaussian random field (GRF) inference on the pTFCE image using the distribution of the global maximum of the test statistic. We also evaluate inference for the pTFCE image using the distribution of the global maximum using the resampling-based procedures. Using the resampling distributions is essentially a semiparametric version of the GRF-based version provided by the package.

## 4 Evaluation methods

### 4.1 Synthetic Simulation Setup

To evaluate the properties of the different resampling inference procedures with and without known heteroskedasticity we performed 1000 simulations of a 1-dimensional image with *n* = 500, *V* = 100, an AR(1) covariance structure with parameter *ρ* = 0.8 with heteroskedasticity across space. We sampled data from 4 groups with no effect on the mean assuming errors were normally distributed. When simulating under homoskedasticity there was no effect of group on the variance of the outcome image, whereas under heteroskedasticity the variances differed across groups (the smallest group variance was 1/20 the size of the largest group variance and it increase linearly across the four groups).

For inference, we compute parametric and robust test statistics and consider two CFTs *p ∈* {0.01, 0.001} for CEI and CMI, as well as inference on local maxima. We use the normal and Rademacher bootstrap, and permutation resampling procedures described in Section 3.1 to estimate the distribution of these topological features. Details and code are given in the Supplementary Material.

### 4.2 Bootstrap-based Simulation Setup

We use 500 bootstrap-based simulations using 682 Voxel-based morphometry (VBM) outcome images from the NKI-RS to evaluate the methods described above in terms of their finite sample performance in two different scenarios for the three different procedures described above. This low number of simulations is presented with 90% probability intervals because of the computational demand of performing 500 bootstraps for each sample size, resampling method, and test statistic value. We consider three sample sizes *n* ∈ {25, 100, 200} and all other simulation settings are as described in Section 4.1 In addition to those procedures, pTFCE is performed using the default settings in the `R` package `pTFCE`. We report the type 1 error rate for the *α* = 0.05 threshold provided by the GRF default in the pTFCE package, and also use the resampled distribution of the global maximum to perform inference for the pTFCE image.

Details of data processing and quality assurance are provided in the Supplement. Briefly, prior to processing, 695 people with demographics and T1 images were visually inspected for quality. A total of 14 scans were excluded because they failed processing or had significant motion or artifacts. VBM was conducted in the Computational Anatomy Toolbox 12 (CAT12: Version 12.5) in Statistical Parametric Mapping 12 (SPM12: Version 7487; Ashburner and Friston, 2005; Ashburner, 2009). After estimation the VBM images were registered to the MNI template and downsampled to 2mm isotropic resolution (Ashburner, 2007). To further speed computation for the simulations, we restrict the study region to 11 axial slices in the center of the image, which included 51,925 voxels.

We use the VBM images to generate realistic data under the null: first, we regress out the effects of covariates in the full sample of 682 subjects with complete scans using the model

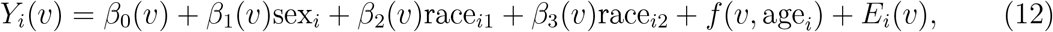

where sex_*i*_ and race_*ij*_ were indicator variables identifying sex and race (Black, Other, White), and *f* was a nonlinear function fit with unpenalized natural cubic splines with 10 degrees of freedom using the ns package in `R`. In each simulation we draw a bootstrap sample 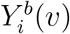 from the image residuals, *E*_*i*_(*v*), and covariates of model (12). Here, we consider one scenario for testing the association between a covariate and the imaging outcome, and two other types of covariates are described and presented in the Supplementary Material. In each bootstrap, we fit the model

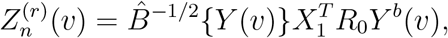

where *f* is estimated with unpenalized natural cubic splines on 4 degrees of freedom and tested it against a model with a linear effect of age yielding a test statistic with 3 degrees of freedom.

Because we residualized the images to the covariates in the full sample, there is no association between the mean of the images in the sample and the covariate. Homoskedasticity is not necessarily satisfied, because only the effect of the covariate on the mean has been removed by the regression (12), but the covariate may still have an effect on the covariance function of the image. The effect of the covariate on the covariance function is implied in fMRI data by studies that demonstrate connectivity differences between diagnosis, age, or motion (for discussion, see Vandekar et al., 2019). In VBM, this can occur when the variance is affected by covariates, such as sex (Forde et al., 2020). Our goal is to understand the how the different resampling methods perform under this realistic setting of heteroskedasticity.

### 4.3 Evaluation metrics

We use two approaches for assessing the accuracy of the resampling methods. First we assess the type 1 error rate at thresholds for *α ∈ {*0.01, 0.05, 0.1, 0.25}. This metric is useful to assess how appropriate the resampling method is for performing hypothesis testing. For the marginal distributions, we apply BH procedure to the uncorrected p-values and compare the estimated and expected type 1 error rate for each threshold. This approach assesses the weak control of the procedure, since the null is true everywhere (Lehmann and Romano, 2005). For the global distributions we report the type 1 error rate of the adjusted p-values found by comparing the minimum adjusted p-value across all observed maxima or clusters in each simulation to the *α* level.

For the second approach we compare the simulated quantiles to those estimated by the resampled distribution. If the resampled distribution perfectly matches the distribution of the simulated test statistic then the quantiles will fall on the identity line.

## 5 Synthetic Simulation Results

We simulated data under known heteroskedasticity to evaluate how the different resampling inference procedures perform in reproducing the distributions of TFs under the null. All resampling methods had near nominal type 1 error rates under homoskedasticity for parametric and robust test statistics, with the permutation and Rademacher bootstrap methods having slightly better accuracy than the normal bootstrap (**Figure 1**). Under heteroskedasticity the permutation procedure had inflated error rates for parametric statistics, but not for robust test statistics. The two bootstrap methods had near nominal type 1 error rates. Simply using a robust covariance estimator resolves the problem with the permutation inference procedure because the procedure is scaling out the differences in the variances across subjects in each permutation by computing the square root covariance matrix as a function of the permuted data (7).

**Figure 1.**
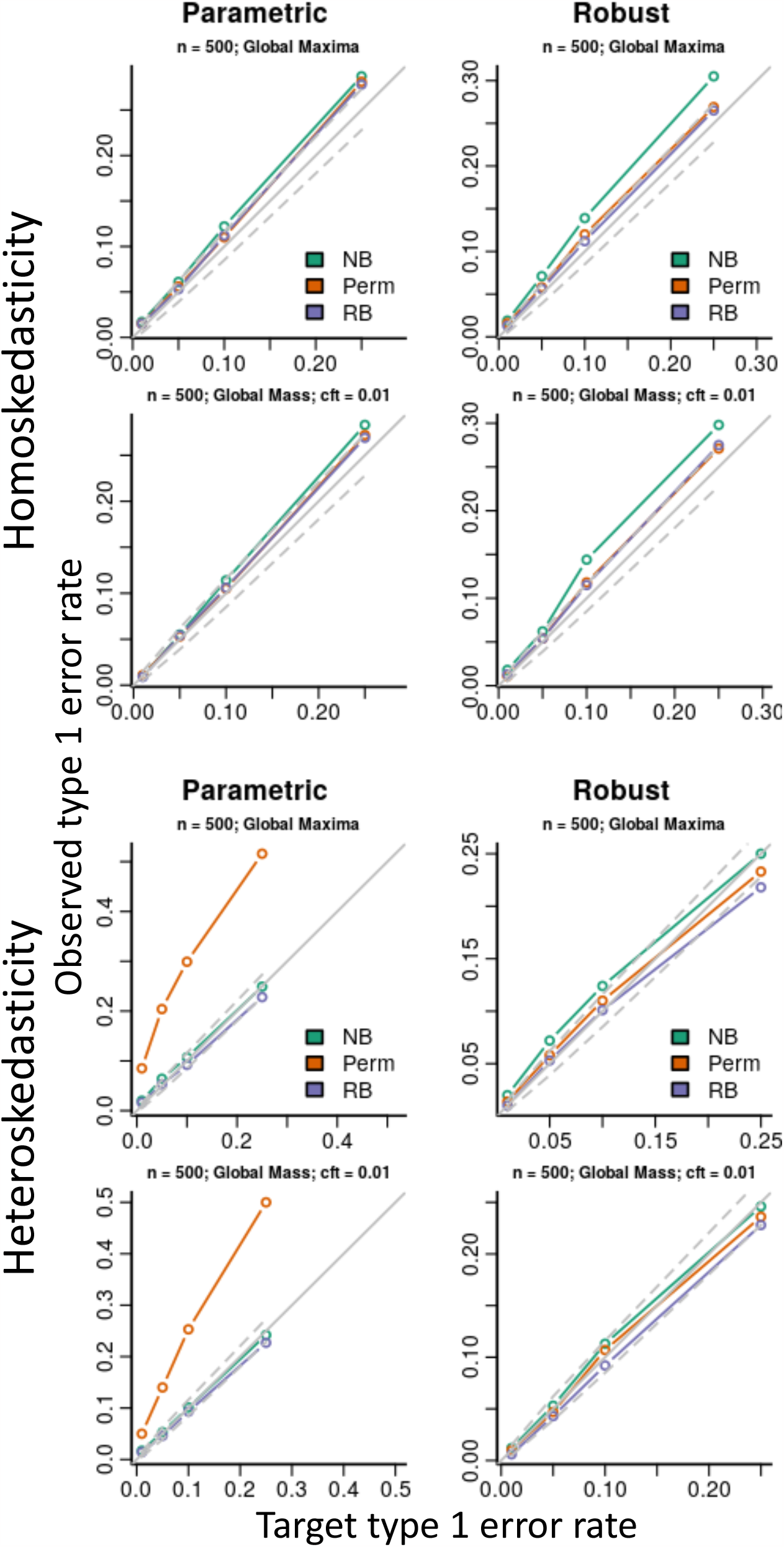
Simulation type 1 error rate results for a 1-dimensional image and sample size of *n* = 500 under homo- and heteroskedasticity (see Section 4.1).

## 6 Bootstrap-based Simulation Results

We assess type 1 error rates for testing global maxima of the parametric and robust statistical images. For the parametric images, all methods perform well in small and large samples across all types of global TF and CFTs considered except the Global pTFCE method, which is conservative across all sample sizes, as indicated by the lines being below the probability interval in the last row (**Figure 2**). For the robust statistical images the normal bootstrap has highly inflated error rates for estimating the distribution of all the topological features, with the exception of the Global pTFCE method (**Figure 3**). The permutation procedure has closer to the nominal error rate in small samples than the two bootstrap procedures, but the Rademacher bootstrap is comparable in samples larger than 50. As for the parametric statistics, these resampling procedures are conservative for the Global pTFCE approach.

**Figure 2.**
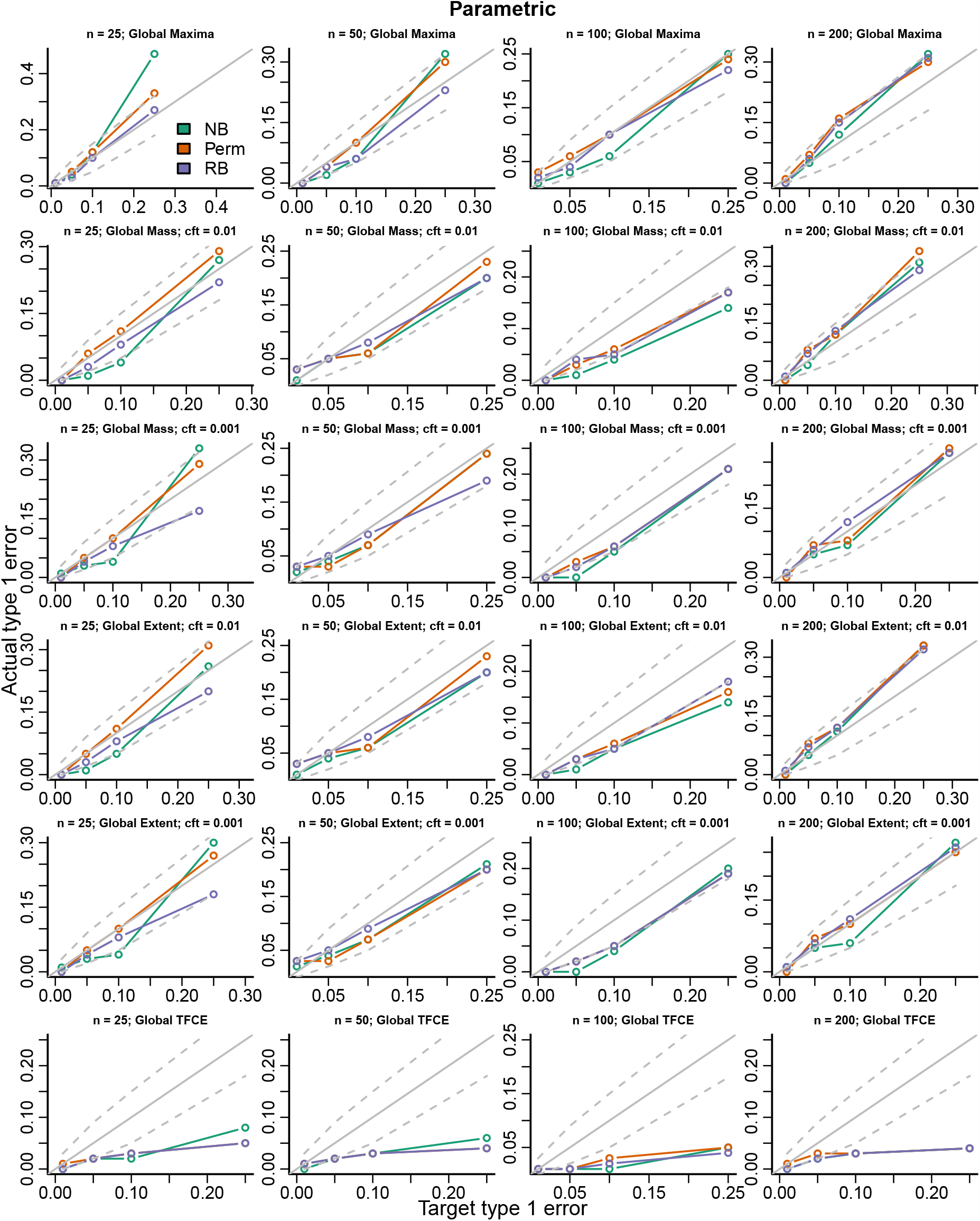
Actual versus target type 1 error rates for testing the global maximum of each topological feature (TF) of the parametric test statistic image. Each row corresponds to a type of TF and each column corresponds to a sample size (Section 3.2). The solid gray line is the nominal type 1 error rate and the dashed gray lines are 90% probability intervals with parameter equal to the target error rate. NB normal bootstrap; Perm permutation; RB Rademacher bootstrap.

**Figure 3.**
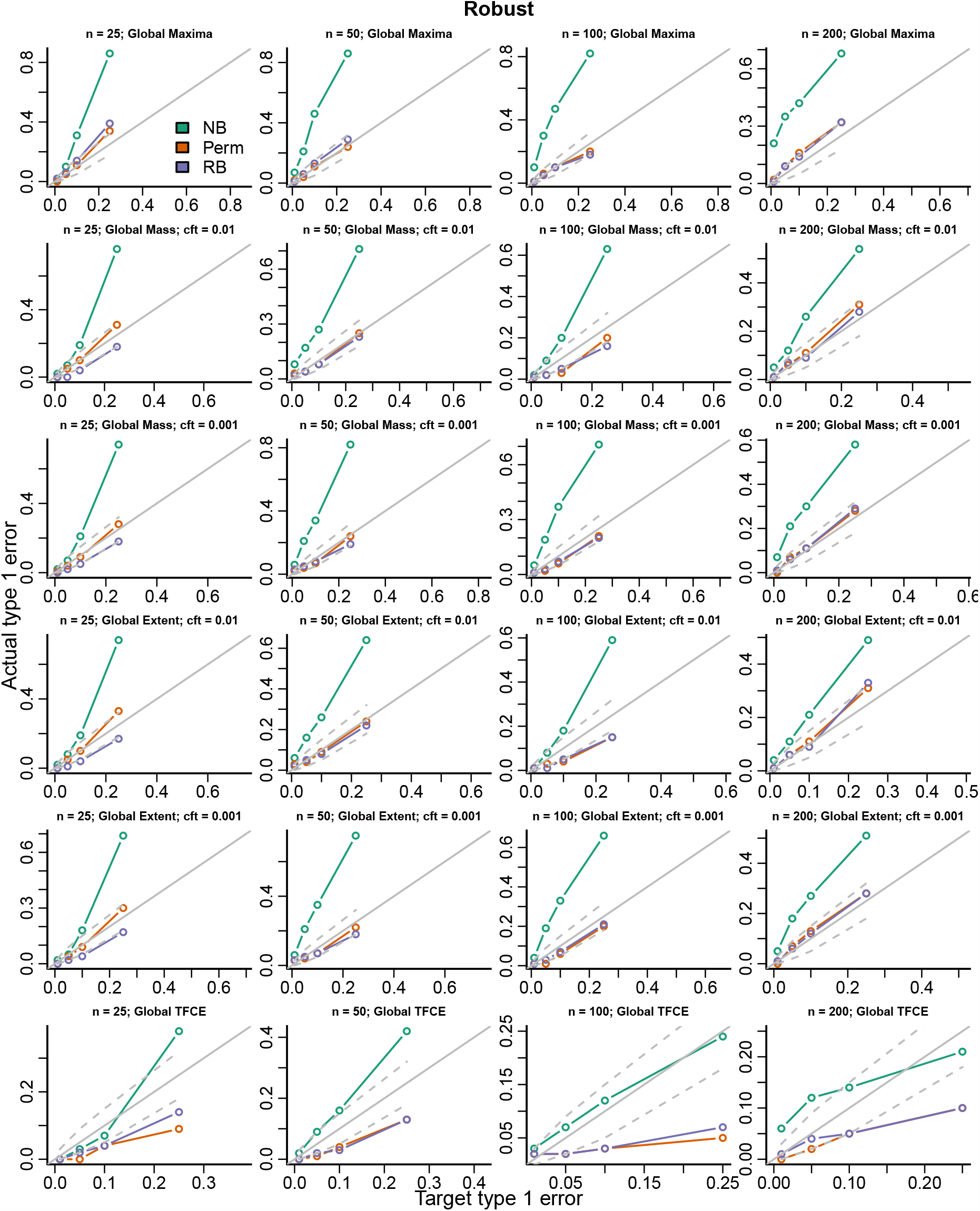
Actual versus target type 1 error rates for testing the global maximum of each topological feature (TF) of the robust test statistic image. Each row corresponds to a type of TF and each column corresponds to a sample size (Section 3.2). The solid gray line is the nominal type 1 error rate and the dashed gray lines are 90% probability intervals with parameter equal to the target error rate. NB normal bootstrap; Perm permutation; RB Rademacher bootstrap.

We simulated data by resampling images, which will retain the heteroskedasticity that exists in the data. The heteroskedasticity that exists in these data does not seem to affect the inference for the age effect, suggesting that permutation testing is robust to the amount of heteroskedasticity that researchers might see in neuroimaging studies like the NKI-RS.

We compared estimated quantiles between the resampled distributions and the simulated distribution to assess how well each procedure reproduces the true distribution. Ideally, the resampled quantiles should match the simulated quantiles. If a method tends to lie above the line it will be conservative (over-estimate quantiles), if it is under the line it will be anti-conservative and underestimate the quantiles. For the parametric statistical image, The permutation sits closer to the gray line indicating better fit, the Rademacher bootstrap is slightly higher in smaller samples, but is otherwise similar to the permutation method for larger samples (**Figure 4**). The normal bootstrap has larger estimated quantiles than the other methods for all sample sizes. For the robust statistical image, the normal bootstrap has worse accuracy than the other two methods; it tends to sit below the line, so is anticonservative. which contributes to the higher type 1 error rates (**Figure 5**). The permutation procedure performs better (is closer to the gray line) than the Rademacher bootstrap across sample sizes, but particularly with the smallest sample size. The plots suggest the Rademacher (and normal) bootstrap has greater spread in the quantiles estimates than the permutation procedure as indicated by purple lines distributed farther away from the gray target line. This is likely related to the fact that, using the bootstrap procedures, the covariates are tied to their outcome images, where as with the permutation procedure the outcome images are permuted. While the permutation assumes exchangeability, if the effect of heteroskedasticity is small for this covariate, then permuting the labels will yield a more stable estimate of the distribution of the TFs.

**Figure 4.**
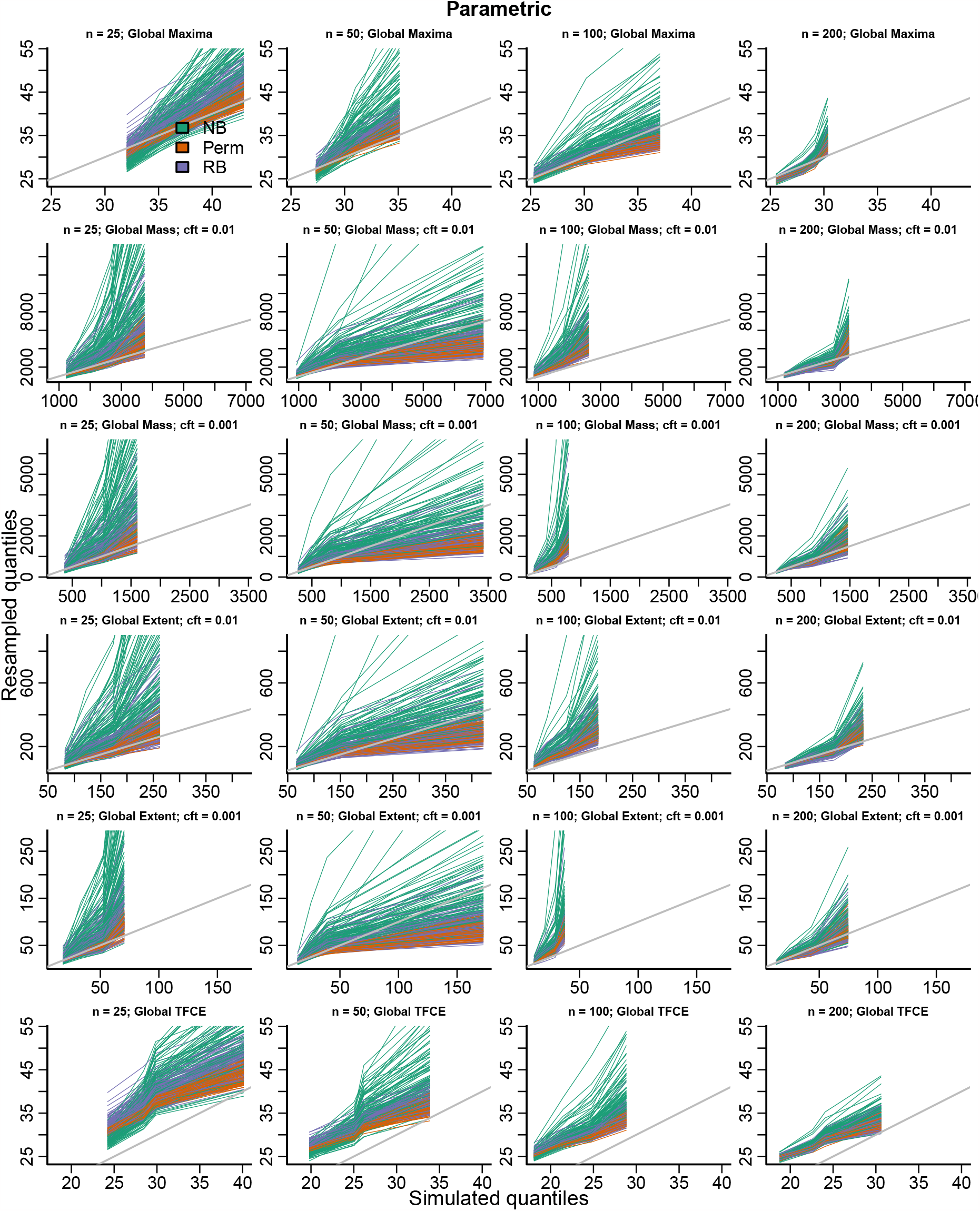
QQ plots comparing resampled distributions to the ground truth simulated distribution for the parametric test statistics. Each colored line corresponds to the quantiles of the simulations (x-axis) versus the quantiles of the resampling procedure in one of the simulations (y-axis). Each row corresponds to a particular type of TF and each column corresponds to a sample size (Section 3.2). The gray line has intercept zero and slope 1. NB normal bootstrap; Perm permutation; RB Rademacher bootstrap.

**Figure 5.**
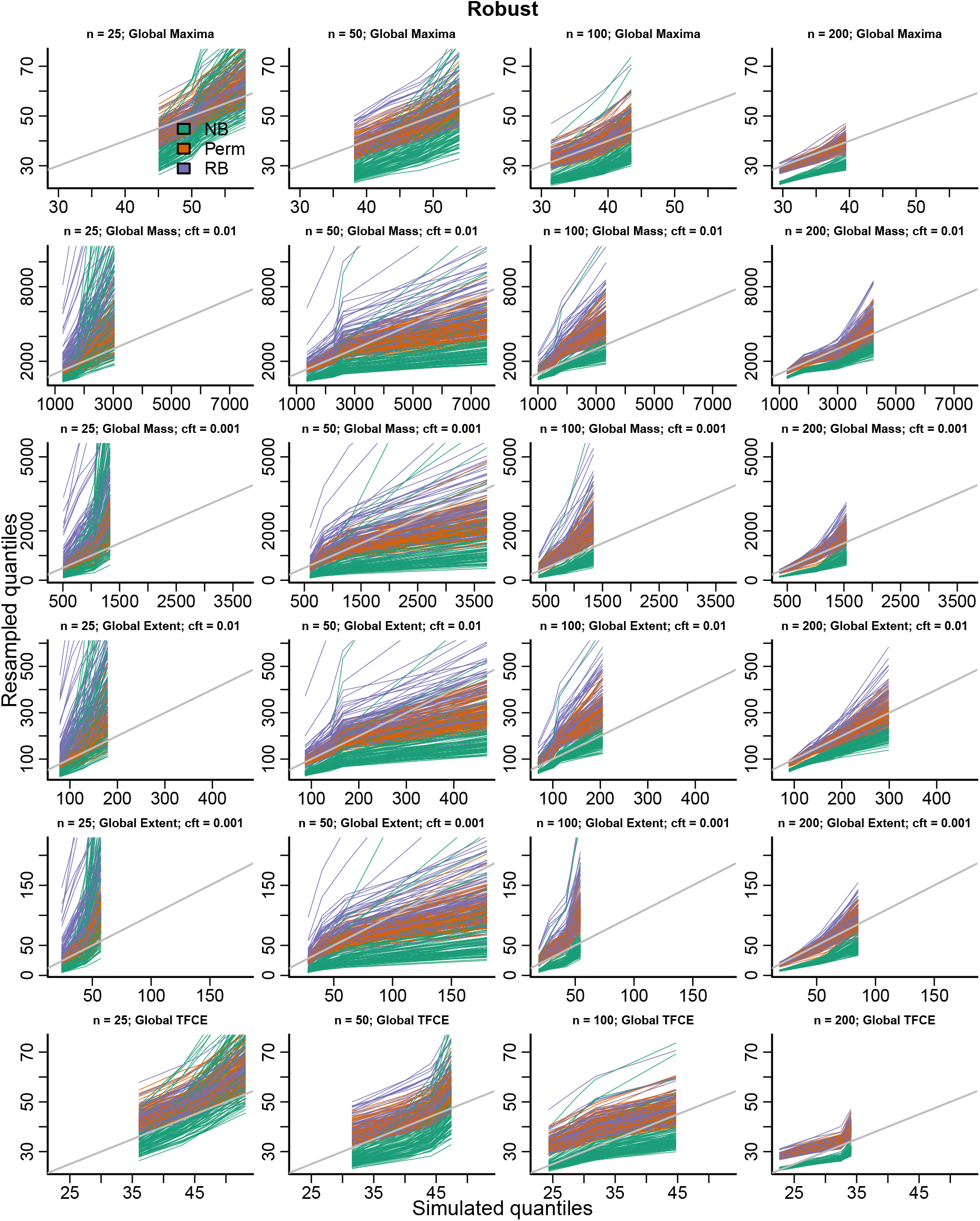
QQ plots comparing resampled distributions to the ground truth simulated distribution for the parametric test statistics. Each colored line corresponds to the quantiles of the simulations (x-axis) versus the quantiles of the resampling procedure in one of the simulations (y-axis). Each row corresponds to a particular type of TF and each column corresponds to a sample size (Section 3.2). The gray line has intercept zero and slope 1. NB normal bootstrap; Perm permutation; RB Rademacher bootstrap.

Across both figures it is possible to notice a pattern in the change of the quantiles of the simulated data on the x-axis for statistics that depend on the value of the test statistic image (Global Maxima, Global Mass, Global pTFCE). For example, in Figure 5, the quantiles shift to the left with increasing sample size. This is due to the convergence of the TF to its asymptotic distribution determined by (S7).

Results for using the GRF-based inference provided in the `pTFCE` package are provided in Table 1. For parametric and robust test statistics GRF-based inference has an inflated type 1 error in small samples that reduces to the nominal level with increasing sample size.

**Table 1:** Type 1 error rates for *α* = 0.05 using GRF-base inference for the pTFCE image.

| n | Parametric | Robust |
| --- | --- | --- |
| 25 | 0.23 | 0.76 |
| 50 | 0.11 | 0.51 |
| 100 | 0.06 | 0.23 |
| 200 | 0.04 | 0.17 |

Estimation for the marginal distributions of all TFs was effective for all sample sizes considered for the Rademacher bootstrap and Permutation (Figures S20 & S20).

In summary, XX.

## 7 Nonlinear age-related anatomical changes in the NKI-RS

Sample characteristics are given in Table 2. In order to study sex differences in the mean adult life-span curve of gray matter volume we fit the model

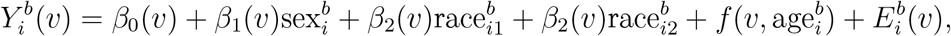

where *f*_1_ and *f*_2_ were fit with natural cubic splines on 4 degrees of freedom. We used parametric and robust test statistic images with each of the the permutation and Rademacher bootstrap methods to test the nonlinear age by sex interaction, which was modeled with 4 degrees of freedom and we performed the analysis using CEI and CMI. This analysis was performed in the full image using the full data set with 5000 bootstrap/permuted samples. The bootstrap and permutation procedures took 13.73 and 13.17 hours to run, respectively. For the parametric statistic the unadjusted p-values computed using (10) were similar between the permutation and bootstrap procedures, however the FWER-adjusted p-values using the distribution of the maximum were substantially different between the Rademacher bootstrap and permutation procedures, with the bootstrap p-values being more conservative (**Table 3**).

**Table 2:**
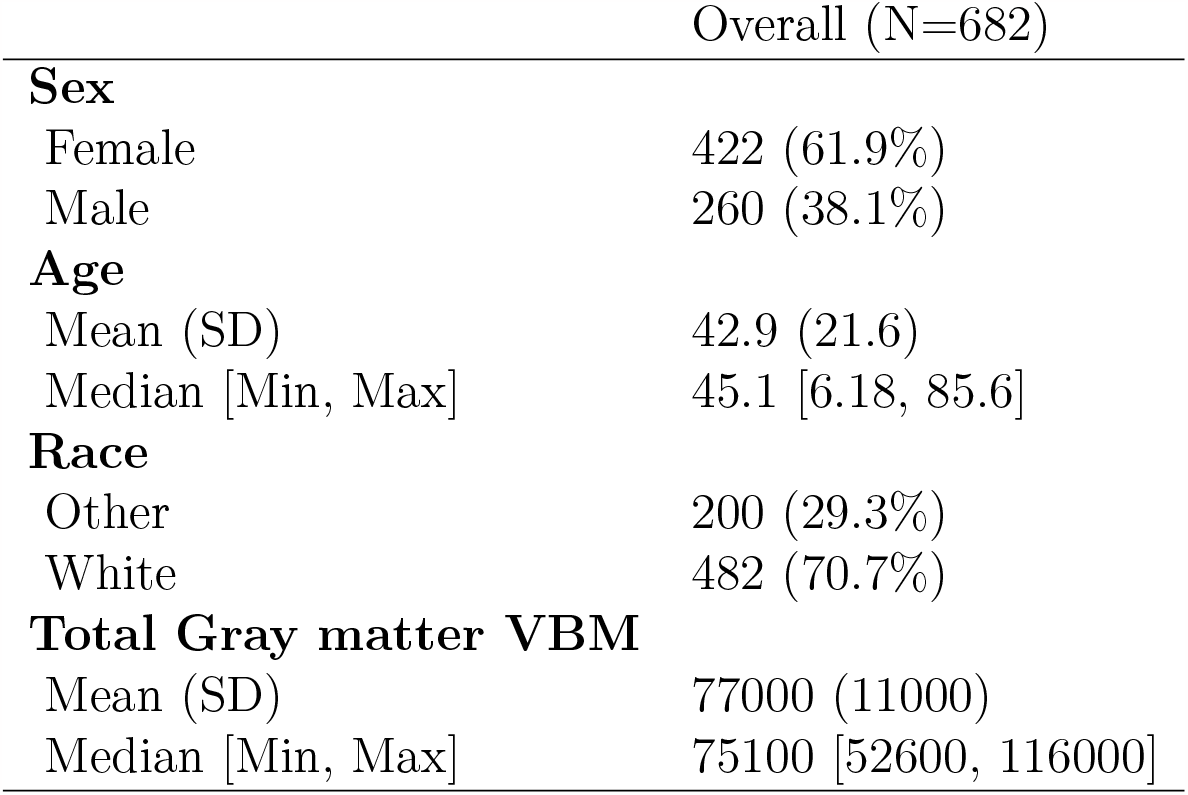
Sample characteristics for the NKI-RS data. Total Gray matter VBM is the sum of the VBM data across each participants image.

**Table 3:** Cluster extent inference p-values for the largest 10 suprathreshold clusters using the parametric test statistic image for the test of the nonlinear age by sex interaction in the NKI-RS data. Cluster extent is in voxels (2mm isotropic resolution) and the cluster centroid is in voxel coordinates. Unadj are unadjusted cluster p-values using formula (10); FWER-adj are adjusted p-values using the distribution of the maximum cluster size; boot and perm correspond to the Rademacher bootstrap and permutation procedure, respectively.

| Cluster Extent | Centroid (vox) | Unadj boot | Unadj perm | FWER-adj boot | FWER-adj perm |
| --- | --- | --- | --- | --- | --- |
| 384 | 46, 88, 28 | 0.000 | 0.000 | 0.108 | 0.059 |
| 288 | 30, 33, 25 | 0.001 | 0.000 | 0.220 | 0.117 |
| 189 | 59, 29, 25 | 0.002 | 0.001 | 0.451 | 0.272 |
| 187 | 36, 26, 13 | 0.002 | 0.001 | 0.463 | 0.274 |
| 173 | 23, 53, 66 | 0.002 | 0.002 | 0.502 | 0.311 |
| 171 | 73, 31, 34 | 0.002 | 0.002 | 0.509 | 0.312 |
| 170 | 42, 59, 42 | 0.002 | 0.002 | 0.514 | 0.314 |
| 122 | 46, 31, 47 | 0.004 | 0.003 | 0.716 | 0.477 |
| 121 | 52, 39, 10 | 0.005 | 0.003 | 0.722 | 0.481 |
| 116 | 39, 55, 74 | 0.005 | 0.003 | 0.754 | 0.493 |

For the robust statistic the unadjusted and FWER-adjusted p-values using (10) were similar across the two procedures (**Table 4**). There was one large cluster in medial frontal cortex that passed the classical *α* = 0.05 adjusted threshold (**Figure 6**). In this region, males and females showed a different pattern of gray matter volume reduction through the life-span. Females declined rapidly until age 35-40, where there was a reduction in the rate of decline through the remainder of the life-span. Males showed a similar pattern of rapid decline, had constant gray matter volume from ages 35-45, and then declined rapidly after age 45. These findings are corroborated by previous studies in development and aging (Gennatas et al., 2017; Chen et al., 2007; Armstrong et al., 2019). The findings are likely to represent, at least in part, differences in curve of total intracranial volume between males and females, but also characterize potential differences in the lifespan course of gray matter volume in orbital gyrus.

**Table 4:** Cluster extent inference p-values for the largest 10 suprathreshold clusters using the robust test statistic image for the test of the nonlinear age by sex interaction in the NKI-RS data. Cluster extent is in voxels (2mm isotropic resolution) and the cluster centroid is in voxel coordinates. Unadj are unadjusted cluster p-values using formula (10); FWER-adj are adjusted p-values using the distribution of the maximum cluster size; boot and perm correspond to the Rademacher bootstrap and permutation procedure, respectively.

| Cluster Extent | Centroid (vox) | Unadj boot | Unadj perm | FWER-adj boot | FWER-adj perm |
| --- | --- | --- | --- | --- | --- |
| 405 | 45, 89, 28 | 0.000 | 0.000 | 0.048 | 0.045 |
| 210 | 29, 34, 25 | 0.001 | 0.001 | 0.214 | 0.199 |
| 155 | 41, 58, 42 | 0.002 | 0.002 | 0.354 | 0.342 |
| 139 | 52, 40, 10 | 0.002 | 0.002 | 0.397 | 0.407 |
| 134 | 73, 31, 34 | 0.002 | 0.002 | 0.417 | 0.434 |
| 115 | 36, 27, 13 | 0.003 | 0.003 | 0.508 | 0.534 |
| 110 | 59, 29, 25 | 0.003 | 0.003 | 0.541 | 0.557 |
| 101 | 71, 63, 36 | 0.004 | 0.004 | 0.596 | 0.607 |
| 100 | 23, 53, 67 | 0.004 | 0.004 | 0.602 | 0.612 |
| 96 | 47, 32, 47 | 0.004 | 0.004 | 0.621 | 0.641 |

**Figure 6.**
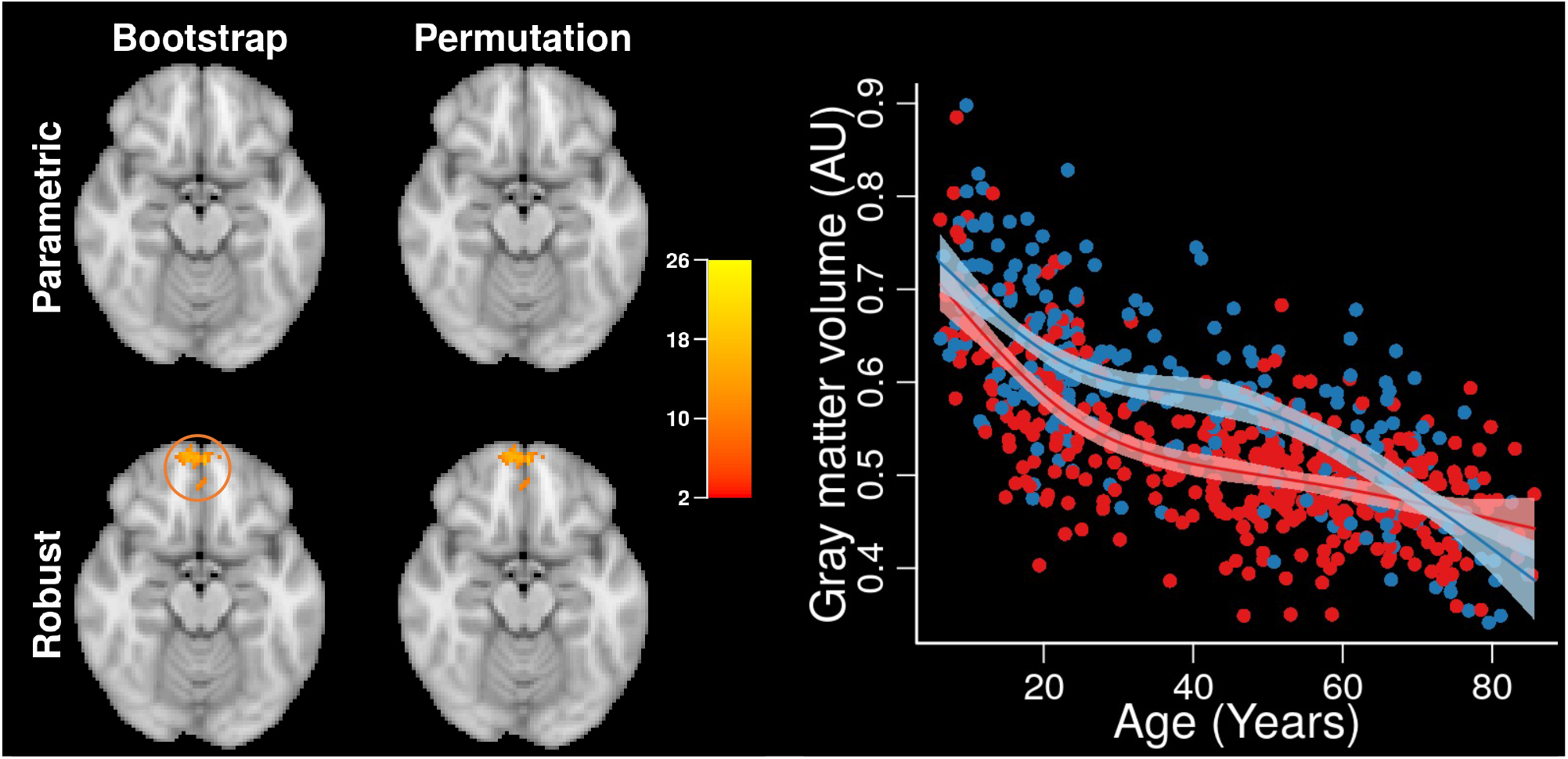
Cluster-extent inference results for the test of the age by sex interaction in the NKI-RS data at axial slice *Z* = 28. There was evidence for difference in the life-span trajectory in the medial frontal cortex. The image shows clusters with *p* < 0.05 for the parametric and robust test statistic images (rows) for the permutation and Rademacher bootstrap procedures (columns). The red-yellow image intensity is −log_10_(*p*) uncorrected p-values. The plot shows the mean trajectory for males (blue) and females (red) within the cluster detected by the robust test statistic image.

## 8 Discussion

Here, we compared three resampling-based procedures for estimating the null distribution of TF of the test statistic image. Our simulation results suggest the commonly used permutation procedure and the Rademacher wild bootstrap are most effective at estimating the null distribution. The bootstrap has the advantage that it weakens the exchangeability assumption required by the permutation procedure, but assumes that the errors are symmetric. Accommodating nonexchangeability is important in neuroimaging studies where it is likely that the variance or covariance of the outcome image may be associated with covariates (Forde et al., 2020; Ritchie et al., 2018). The permutation approach may have better accuracy and lower variance across samples when the assumption of exchangeability is roughly satisfied and permuting the residuals yields a more stable estimate of the null distribution. In our prevous research, we found that the permutation procedure has some robustness to violations of exchangeability, so appears to be a reliable choice for inference in neuroimaging (Vandekar et al., 2019; Lehmann and Romano, 2005).

We also evaluated the use of the GRF-based pTFCE method and showed that it has higher than the nominal error rate for samples less than 200 for parametric test statistic images and higher than nominal error rate for all samples considered in this study for the robust statistical images. Using the distribution of the maximum estimated using one of the three resampling procedures yields conservative inference and could be used for thresholding pTFCE images in smaller samples instead of the GRF-based threshold.

We evaluated estimation of marginal distributions of TFs, which have been used to compute unadjusted p-values for the TFs in the observed image and then applied with FDR correction. We show that using the Rademacher bootstrap or Permutation procedures controls the weak type 1 error rate (under the global null) for all sample sizes considered. The improved performance may be due to the larger number of features available by considering the marginal distribution (all TFs from each bootstrap are used instead of just the global maximum). In any case, these results suggest this might be a good approach to use in small sample inference.

Relative to existing methods for estimating the distribution of TF of the test statistic images, the resampling approaches have fewer assumptions and are generally applicable regardless of data type, image smoothing, cluster forming threshold, and other processing parameters. These benefits come at the cost of computing time, though computing time can be significantly reduced through the use of parallel computing. Computing time can also be substantially reduced for the permutation procedure using analytic strategies (Winkler et al., 2016) and a similar approach could be possible to speed computation for the bootstrap as well.

Our approach has important limitations: to generate realistic simulated data, we boot-strapped from a large sample of images from the NKI-RS, however, this sample is too small to estimate a full rank covariance matrix for the imaging data. More sophisticated sampling procedures should be developed for high-dimensional data to sample realistic data while appropriately modeling variability in the distribution. In addition, we only focused on estimation under the null, and estimation of distributions of TF under the alternative is likely to be more nuanced (Bowring et al., 2021). Future work can investigate estimation of the distributions of topological features under the alternative and assess coverage rates of confidence sets.

## 9 Funding Acknowledgments

NIH grants R01MH123563 (to S.N.V), R01MH115000 and R01MH102266 (to N.D.W.), R01EB026859 (to A.S).

