## Supplement for "Evaluation of resampling-based inference for topological features of neuroimages"

#### S0.1 Theory

We describe each of the resampling methods in terms a vector of approximately standard normal random variables, $Z_n(v)$ that are a function of the parameter estimators and their variance estimators, and whose sum of squares is proportional to the test statistic image

$$T_n(v) = n \times Z_n^T(v) Z_n^T(v). \tag{S1}$$

This notation allows us to easily describe how each of the resampling procedures computes $Z_n(v)$ using the outcome images $Y(v)$, which gives us the joint distribution of the test statistic image $T_n(v)$.

We use

$$R_0 = I - X_0\{X_0^T X_0\}^{-1} X_0^T$$
$$R = I - X\{X^T X\}^{-1} X^T$$

to denote the symmetric $n \times n$ residual forming matrices. The least squares estimator for the parameter image $\beta(v)$ in (1) can be written as,

$$\hat{\beta}(v) = \{X_1^T R_0 X_1\}^{-1} X_1^T R_0 Y(v).$$

The robust (heteroskedastic consistent) estimator for the covariance is derived from the least squares estimating equation

$$\Psi(\zeta; Y, X) = -n^{-1} \sum_{i=1}^{n} (Y_i(v) - X_i \zeta(v))^2, \tag{S2}$$

which is maximized with respect to $\zeta(v)$ to obtain the estimator $\hat{\zeta}(v)$ (Van der Vaart, 2000). The derivative is equal to zero where the maximum occurs. By using a Taylor expansion of the derivative of (S2) centered at the true value of the parameter, $\zeta(v)$, we obtain

$$\begin{aligned} 0 = \frac{\partial}{\partial \zeta^*} \Psi(\zeta^*; Y, X)\big|_{\hat{\zeta}(v)} &= n^{-1} \sum_{i=1}^{n} \{Y_i(v) - X_i \hat{\zeta}(v)\} X_i^T \\ &= n^{-1} \sum_{i=1}^{n} \{Y_i(v) - X_i \zeta(v)\} X_i^T + -n^{-1} \sum_{i=1}^{n} X_i^T X_i \{\hat{\zeta}(v) - \zeta(v)\} \end{aligned} \tag{S3}$$

where the Frechet derivative is used when differentiating with respect to the function $\zeta^*$ (Van der Vaart, 2000). Note that $X_i$ is a row vector as described in Section 2. Rearranging (S3) to solve for $\sqrt{n}\{\hat{\zeta}(v) - \zeta(v)\}$ gives

$$\begin{aligned} \sqrt{n}\{\hat{\zeta}(v) - \zeta(v)\} &= \left[ n^{-1} \sum_{i=1}^{n} X_i^T X_i \right]^{-1} \times \left[ n^{-1} \sum_{i=1}^{n} \{Y_i(v) - X_i \zeta(v)\} X_i^T \right] \\ &\approx_D \left[ \lim_{n \to \infty} n^{-1} \sum_{i=1}^{n} \mathbb{E} X_i^T X_i \right]^{-1} \times \left[ n^{-1/2} \sum_{i=1}^{n} \{Y_i(v) - X_i \zeta(v)\} X_i^T \right], \end{aligned} \tag{S4}$$

where $\approx_D$ denotes approximately equal in distribution and the second line follows from continuous mapping theorem and Skutsky's theorem. The approximation assumes large sample size. For voxel locations $v, w$, the asymptotic covariance of $\sqrt{n}\{\hat{\zeta}(v) - \zeta(v)\}$ and $\sqrt{n}\{\hat{\zeta}(w) - \zeta(w)\}$ is determined by taking the covariance of the right hand side of (S4). Let $A_\zeta = \lim_{n\to\infty} n^{-1} \sum_{i=1}^{n} \mathbb{E} X_i^T X_i$ and

$$\begin{aligned} B_\zeta(v, w) &= \lim_{n \to \infty} \mathrm{Cov} \left\{ n^{-1/2} \sum_{i=1}^{n} \{Y_i(v) - X_i \zeta(v)\} X_i^T, n^{-1/2} \sum_{i=1}^{n} \{Y_i(w) - X_i \zeta(w)\} X_i^T \right\} \\ &= \lim_{n \to \infty} n^{-1} \sum_{i=1}^{n} \mathbb{E}\{Y_i(v) - X_i \zeta(v)\}\{Y_i(w) - X_i \zeta(w)\} X_i^T X_i, \end{aligned}$$

where the second line follows by independence of the sampling units. Then the asymptotic covariance of $\sqrt{n}\{\hat{\zeta}(v) - \zeta(v)\}$ is

$$\mathrm{Cov}\left\{\sqrt{n}\{\hat{\zeta}(v) - \zeta(v)\}, \sqrt{n}\{\hat{\zeta}(w) - \zeta(w)\}\right\} = A_\zeta^{-1} B_\zeta(v, w) A_\zeta^{-1}. \tag{S5}$$

Because we are only interested in the covariance of the target parameter estimators, $\hat{\beta}(v)$, we want the block diagonal of (S5) that corresponds to $\beta(v)$; the formula is given in Section S6 of the supplement of Vandekar et al. (2019).

The robust (heteroskedastic consistent) estimator for the covariance of $n^{1/2}\{\hat{\beta}(v) - \beta(v)\}$ is a function of

$$\begin{aligned} \hat{A} &= X_1^T R_0 X_1 \\ \hat{B}(v, w) &= X_1^T R_0 \mathrm{diag}\left\{\frac{[RY(v)]_i [RY(w)]_i}{R_{ii} R_{ii}}\right\} R_0 X_1^T, \end{aligned} \tag{S6}$$

where the diag{} operator is a function of the index $i$ that returns an $n \times n$ matrix with the $i$th value on the $i$th diagonal and zeros elsewhere. $B(v, w)$ is a matrix valued functions from the crossproduct of the image domain to $\mathbb{R}^{m_1 \times m_1}$. We showed, in the supplement of Vandekar et al. (2019), that the estimator $A^{-1} B(v, w) A^{-1}$ is consistent for the covariance of $n^{1/2}\{\hat{\beta}(v) = \beta(v)\}$. For a given voxel, $v$, the asymptotic covariance function of the image $n^{1/2}\{\hat{\beta}(v) = \beta(v)\}$ is,

$$\begin{aligned} \Sigma_\beta(v; \theta) &= \mathrm{Cov}\left\{n^{1/2}(\hat{\beta}(v) - \beta_0(v)), n^{1/2}(\hat{\beta}(w) - \beta_0(w))\right\} \\ &= A^{-1} B(v, w) A^{-1}, \end{aligned}$$

where

$$\begin{aligned} A &= \lim_{n\to\infty} n^{-1} \mathbb{E} \hat{A} \\ B(v, w) &= \lim_{n\to\infty} n^{-1} \mathbb{E} X_1^T R_0 \mathrm{diag}\left\{\mathrm{Cov}(Y_i(v), Y_i(w) \mid X_i)\right\} R_0 X_1, \end{aligned}$$

and the expectation is taken over the joint distribution of the data. Under equal variance, $B(v, w)$ simplifies to $A \times \mathrm{Cov}(Y(v), Y(w) \mid X)$. If the weights and the design matrix are identical at each location, then $A$ does not depend on location, $v$, and the formula simplifies to

$$A = \lim_{n\to\infty} n^{-1} \mathbb{E} X_1^T R_0 X_1$$

$A$ does not depend on the location, but $B$ still does through the dependence on the outcome image $Y(v)$. This is the condition we consider in the body of the paper and use in our software. Assuming $A$ does not depend on the location, it is useful to think if $B(v, w)$ as being a function of the image $Y(v)$ instead, which we do using the notation $B\{Y(v), Y(w)\}$.

For a given $v$, there are many proposed estimators for $\hat{B}(v)$ (Long and Ervin, 2000). We rely on the consistent estimator (S6) Long and Ervin (2000) as an approximation to the jackknife version of the sandwich covariance estimator (Huber, 1964). In imaging, we (and others in univariate applications) have found that this estimator works well in finite samples (Vandekar et al., 2019; Long and Ervin, 2000).

We consider two possible versions of (S1), the first is a parametric estimator, that assumes equal variances and covariances across subjects at each location in the image (Vandekar et al., 2019):

$$Z_n(v) = \hat{\sigma}^{-1}(v)\hat{A}^{-1/2}X_1^T R_0 Y(v),$$

where $R_0$ is the residual forming matrix for the full model. The second is a robust estimator (Huber, 1964)

$$Z_n^{\mathrm{Rob}}(v) := \hat{B}^{-1/2}(v,v)\hat{A}\{\hat{\beta}(v) - \beta_0(v)\} = \hat{B}^{-1/2}\{Y(v)\}X_1^T R_0 Y(v).$$

Here the sum of squares of $Z_n(v)$ yields the test statistic image, $T_n(v) = Z_n(v)^T Z_n(v)$. The same is true for the test statistic image using the robust covariance estimator with $Z_n^{\mathrm{Rob}}(v)$. When $m_1 = 1$, $Z_n(v)$ is the T-statistic image for the test of (2).

Under the null, the joint distribution of $Z_n(v)$ is approximately

$$Z_n^{\mathrm{Rob}}(v) \sim N\left\{\mathbf{0}, \Sigma(v,w)\right\}, \tag{S7}$$

where $\Sigma(v,w)$ is the covariance of the test statistic image

$$\Sigma(v,w) = B^{-1/2}(v,v)B(v,w)B^{-1/2}(w,w).$$

For all $v$, $\Sigma(v,v) = I \in \mathbb{R}^{m_1 \times m_1}$ – the normalized statistics have variance one and elements of the statistic vector are independent within each location, $\mathrm{Cov}\{Z_{n,j}(v), Z_{n,k}(v)\} = 0$. In general, $\Sigma(v,w)$ is not diagonal. This implies that the off-diagonal elements of $\Sigma(v,w)$ can be nonzero, i.e. $\mathrm{Cov}\{Z_{n,j}^{\mathrm{Rob}}(v), Z_{n,k}^{\mathrm{Rob}}(w)\} \neq 0$ for $v \neq w$. When, for each $v$, $Y(v)$ is homoskedastic across subjects, then $B(v,w) = \mathrm{Cov}\{Y_i(v), Y_i(w) \mid X_i\}A$. The test statistic image (S1) is asymptotically chi-square at each location, under the null. In our previous paper we were not able to succinctly describe the joint distribution of the image for $m_1 > 1$ (Vandekar et al., 2019). Using the notation we introduce here, we can now express it easily in terms of the distribution of $Z_n(v)$.

### S1 Voxel-based morphometry processing and quality control

Prior to processing, 695 T1 images were visually inspected for quality. Voxel-based morphometry (VBM) completed in 693 scans using the Computational Anatomy Toolbox 12

(CAT12: Version 12.5) in Statistical Parametric Mapping 12 (SPM12: Version 7487; Ashburner and Friston, 2005; Ashburner, 2009). T1-weighted structural images were corrected for bias-field inhomogeneities, registered using linear (12 parameter affine) and non-linear transformations, then spatially normalized using the DARTEL algorithm, and segmented into gray matter, white matter and cerebrospinal fluid (Ashburner, 2007). Further quality control was conducted using the CAT12's quality assurance framework. This is a retrospective, quantitative measure that evaluates image parameters such as noise, inhomogeneities and image resolution, placing images on a rating scale and assigning a letter grade to each image. Any scan rated B- or above is considered good quality, and any scans rated C- to C+ are considered satisfactory. In our data, all CAT12 output rated C+ or below were further visually inspected for gray matter segmentation quality. Any scan with gray matter segmentation that did not capture the gray matter, or included non-gray matter voxels, were excluded from further analysis. A total of 682 participants passed this quality inspection criteria and had complete demographic data.

### S2 Additional simulations

We considered two other scenarios for simulation analyses: scenario 1) we fit the model

$$Y_i^b(v) = \alpha_0 + \alpha_1 \text{sex}_i^b + \alpha_2 \text{race}_{i1}^b + \alpha_2 \text{race}_{i2}^b + \alpha_3 \text{age}_i^b + \alpha_4 X_i^b + \sum_{k=2}^{3} \beta_1 \times \text{sex}_i + E_i^b(v)$$

where $X_i$ is a binary sex covariate; scenario 2) we fit the model

$$Y_i^b(v) = \alpha_0 + \alpha_1 \text{sex}_i^b + \alpha_2 \text{race}_{i1}^b + \alpha_2 \text{race}_{i2}^b + \alpha_3 \text{age}_i^b + \sum_{k=1}^{3} \beta_k I(X_i^b == k) + E_i^b(v),$$

where $X_i$ is a 4 level factor generated nonrandomly and independent of the bootstrap data. The test for each of these scenarios is with 1, and 3 degrees of freedom, respectively. These simulations (where the covariate was synthetically generated) satisfy homoskedasticity because the distribution of the imaging data are independent of the covariate.

### S3 Simulation Results

Details of the simulation analyses and evaluation metrics are given in Section 4 of the paper. This section presents results for all simulations for parametric and robust test statistics evaluating the marginal and global CDFs.

#### S3.1 Sex covariate

The effect of sex was removed from the imaging data using equation (13), and then bootstrap samples of the residuals were modeled as a function of sex and tested on one degree of freedom. Type 1 error rates and quantile-quantile plots comparing the simulated and resampled quantiles are given below.

##### S3.1.1 Marginal distribution

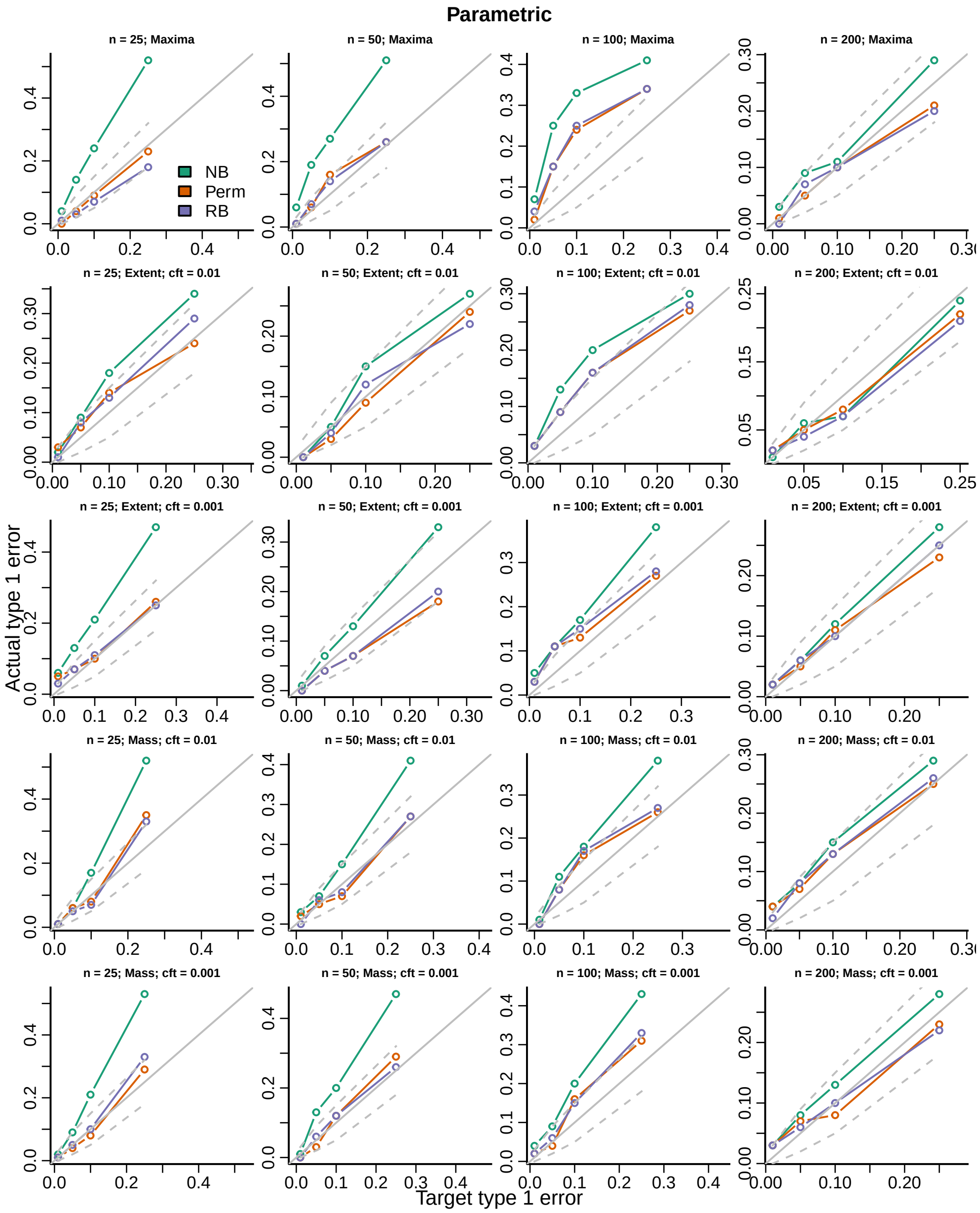


Figure S1: Actual versus target type 1 error rates for the three inference procedures considered for testing the marginal distribution of each topological feature (TF) of the parametric test statistics image.

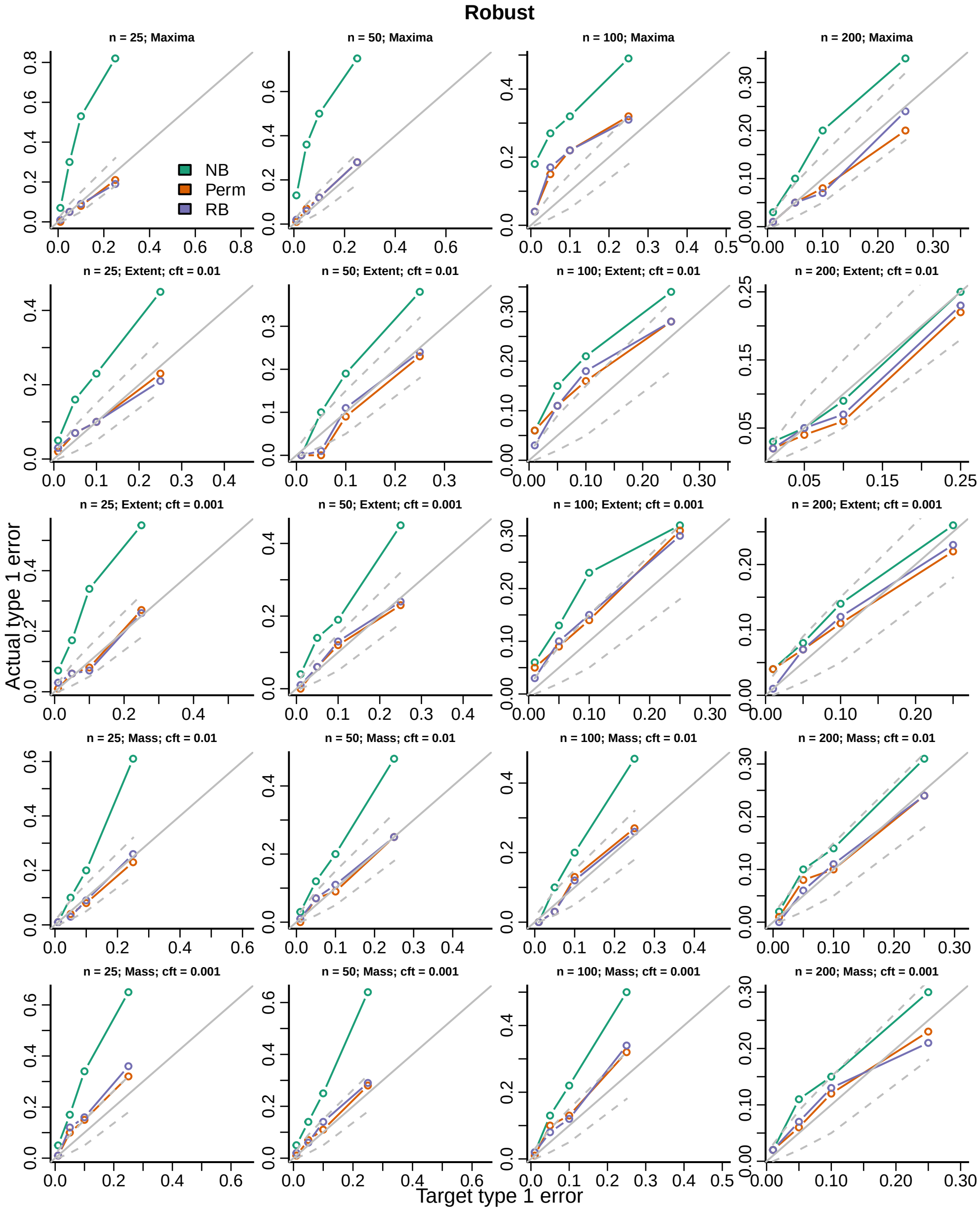


Figure S2: Actual versus target type 1 error rates for the three inference procedures considered for testing the marginal distribution of each topological feature (TF) of the robust test statistics image.

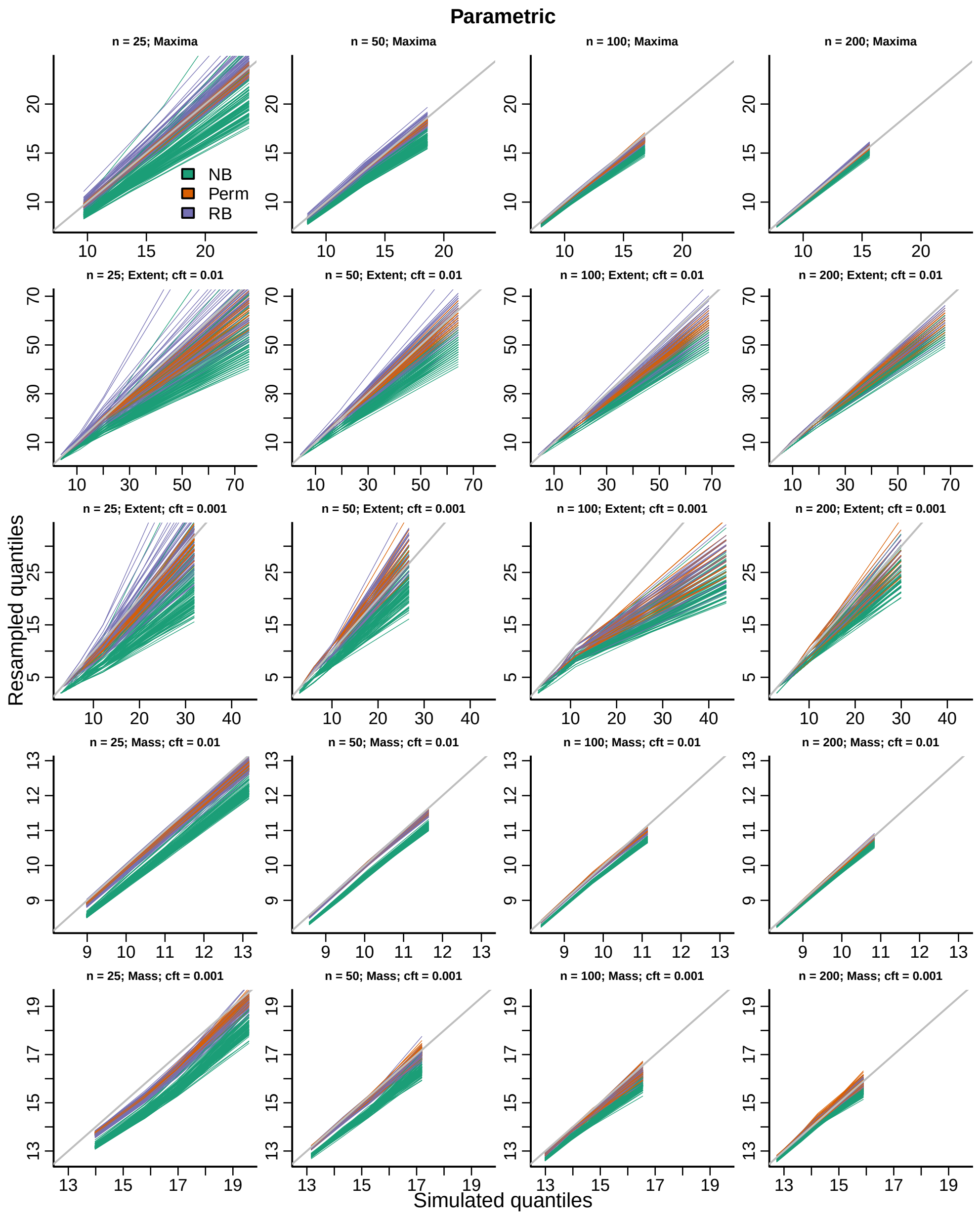


Figure S3: QQ-plots for the three inference procedures considered for the marginal distribution of each topological feature (TF) of the parametric test statistics image.

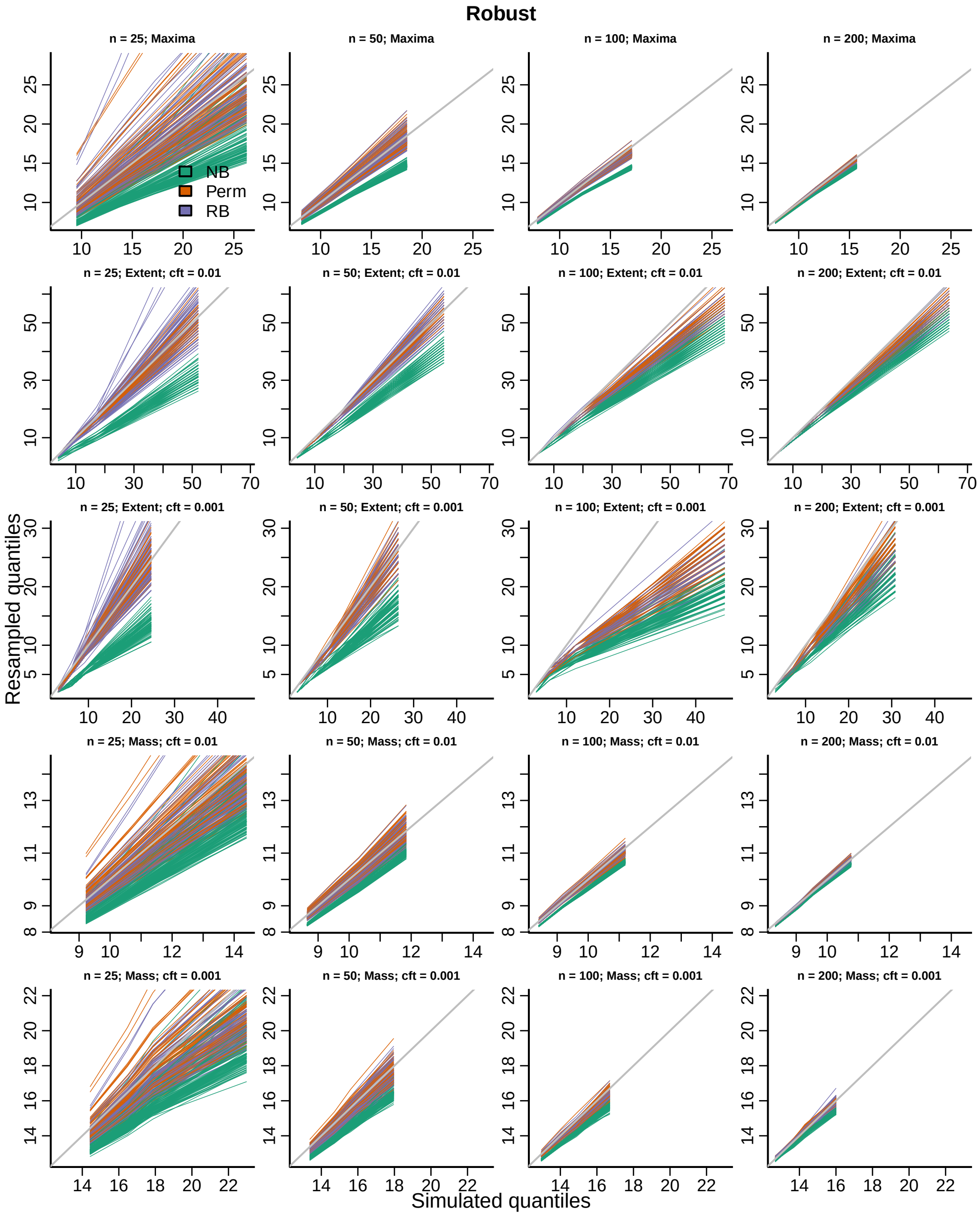


Figure S4: QQ-plots for the three inference procedures considered for the marginal distribution of each topological feature (TF) of the robust test statistics image.

##### S3.1.2 Global distributions

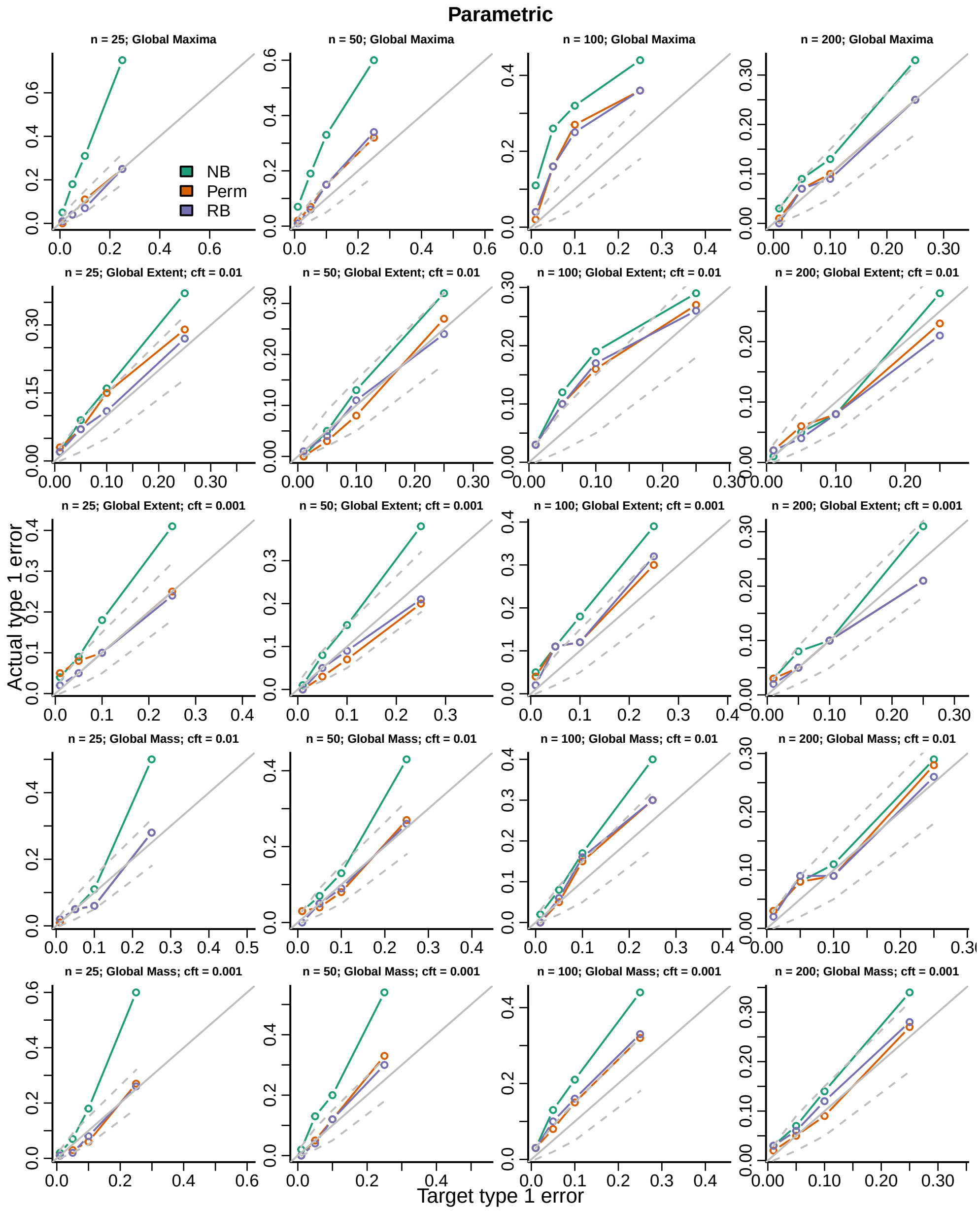


Figure S5: Actual versus target type 1 error rates for the three inference procedures considered for testing the distribution of the global maximum of each topological feature (TF) of the parametric test statistics image.

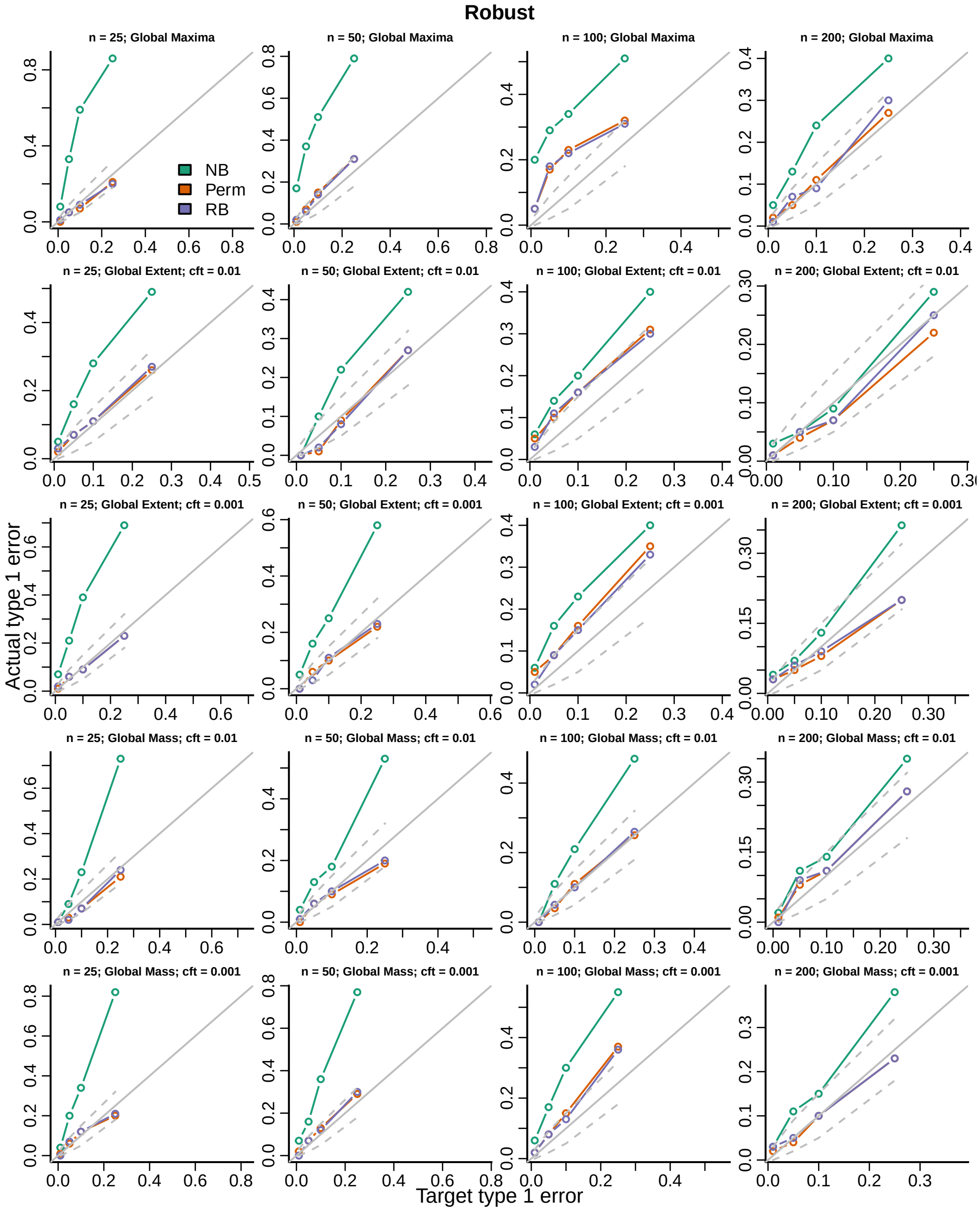


Figure S6: Actual versus target type 1 error rates for the three inference procedures considered for testing the distribution of the global maximum of each topological feature (TF) of the robust test statistics image.

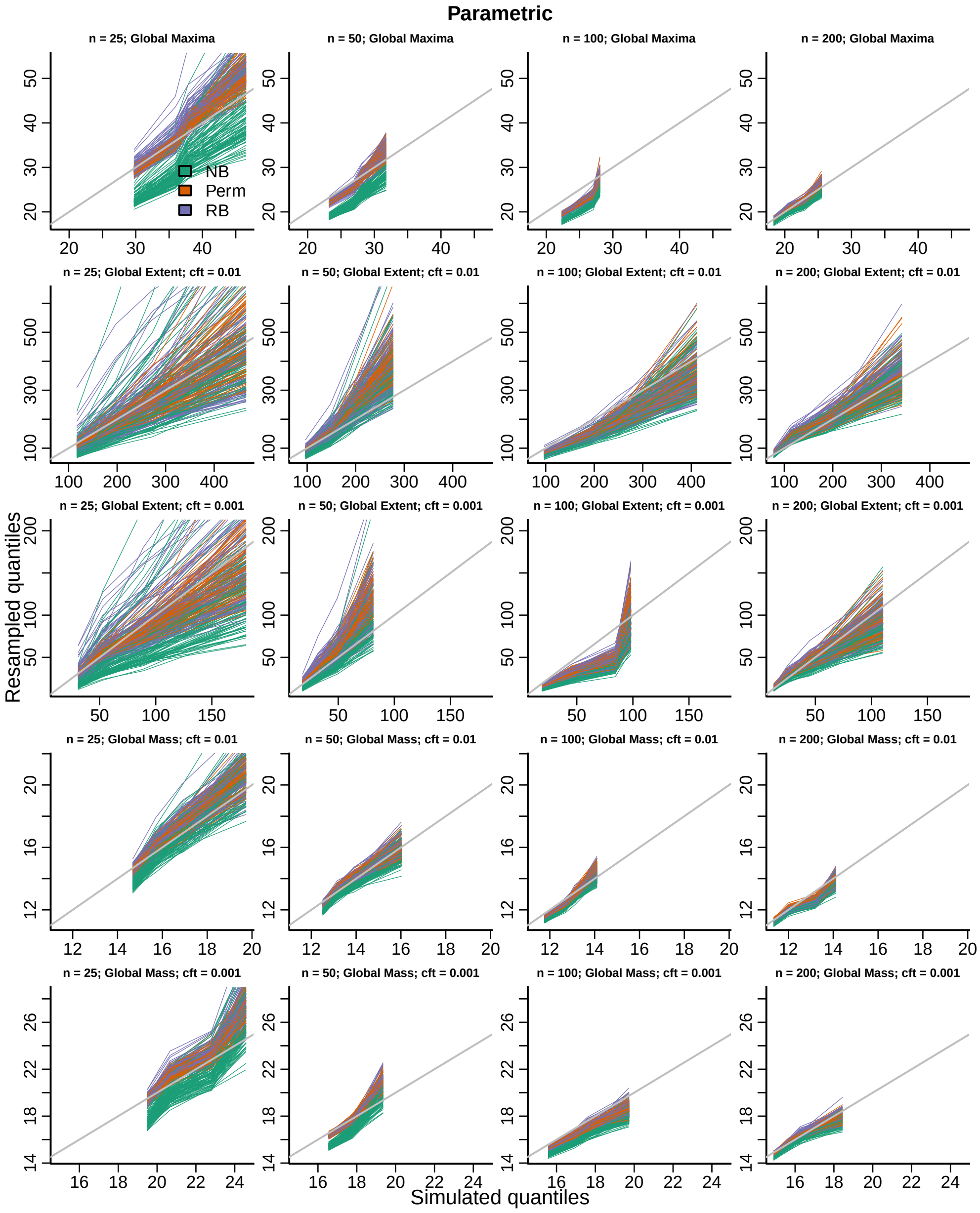


Figure S7: QQ-plots for the three inference procedures considered for the distribution of the global maximum of each topological feature (TF) of the parametric test statistics image.

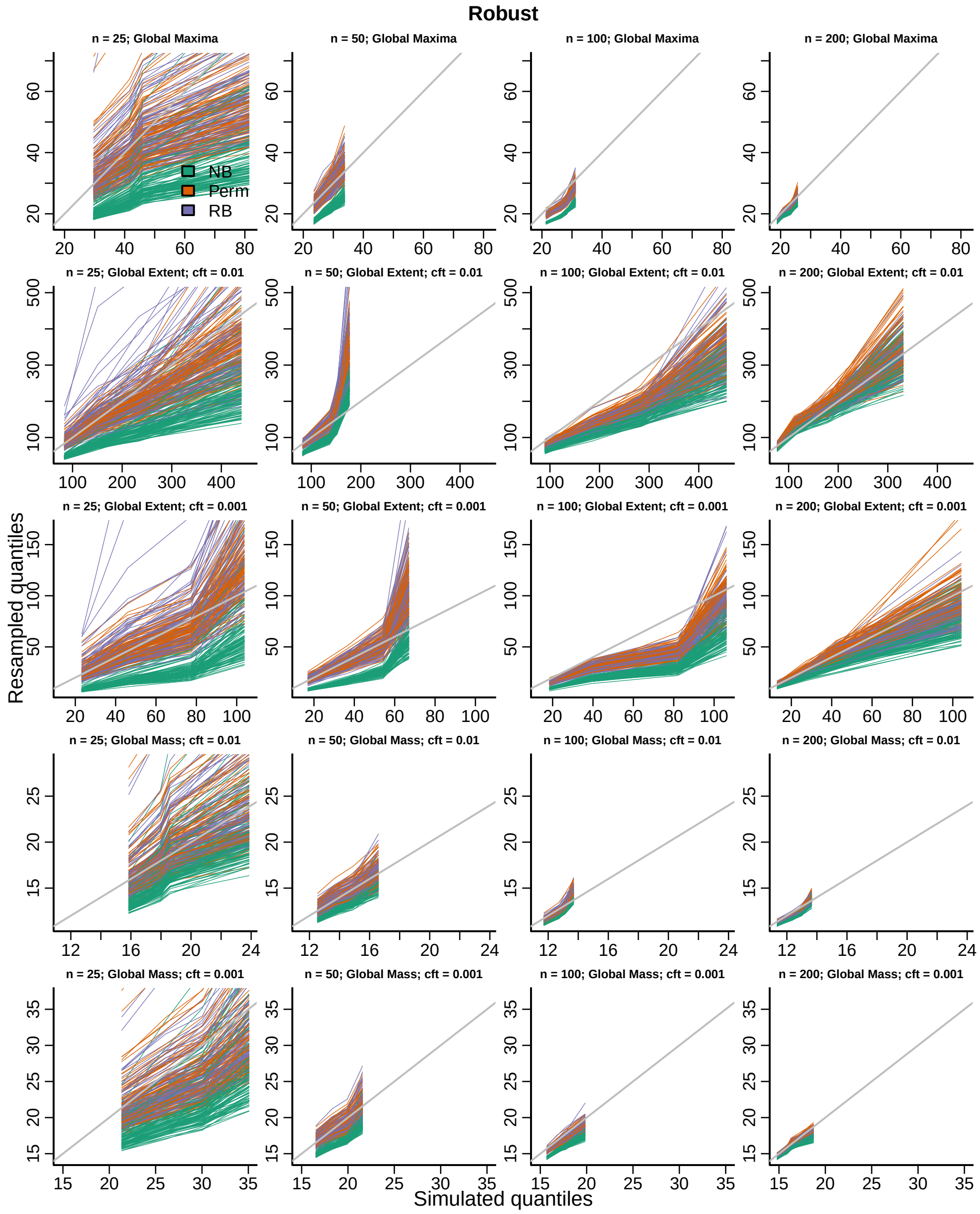


Figure S8: QQ-plots for the three inference procedures considered for the distribution of the global maximum of each topological feature (TF) of the robust test statistics image.

#### S3.2 Fake group variable

Group was simulated independently of the imaging data and the bootstrap samples of the imaging data were modeled and tested on 3 degrees of freedom. Type 1 error rates and QQ-plots are given below.

##### S3.2.1 Marginal distribution

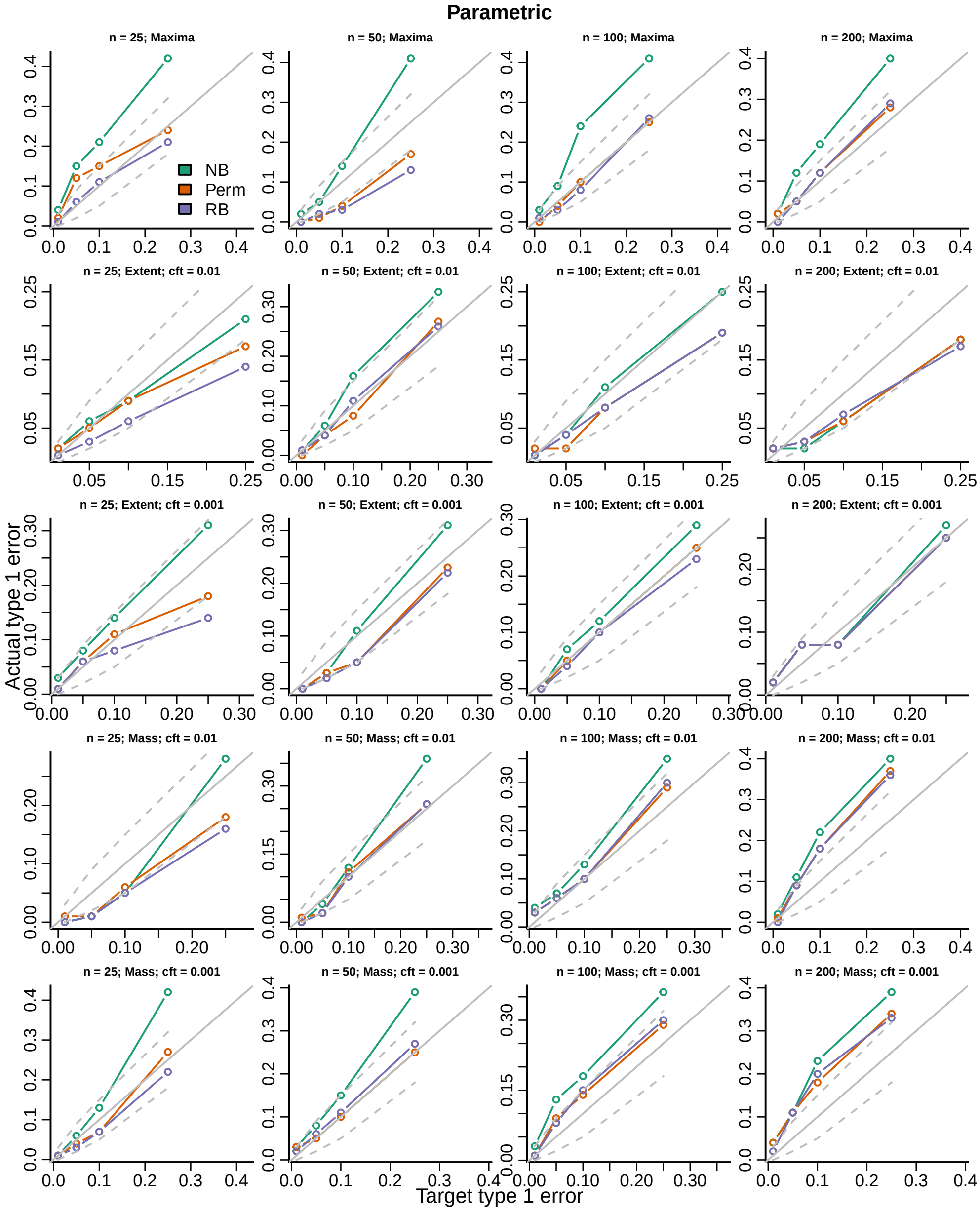


Figure S9: Actual versus target type 1 error rates for the three inference procedures considered for testing the marginal distribution of each topological feature (TF) of the parametric test statistics image.

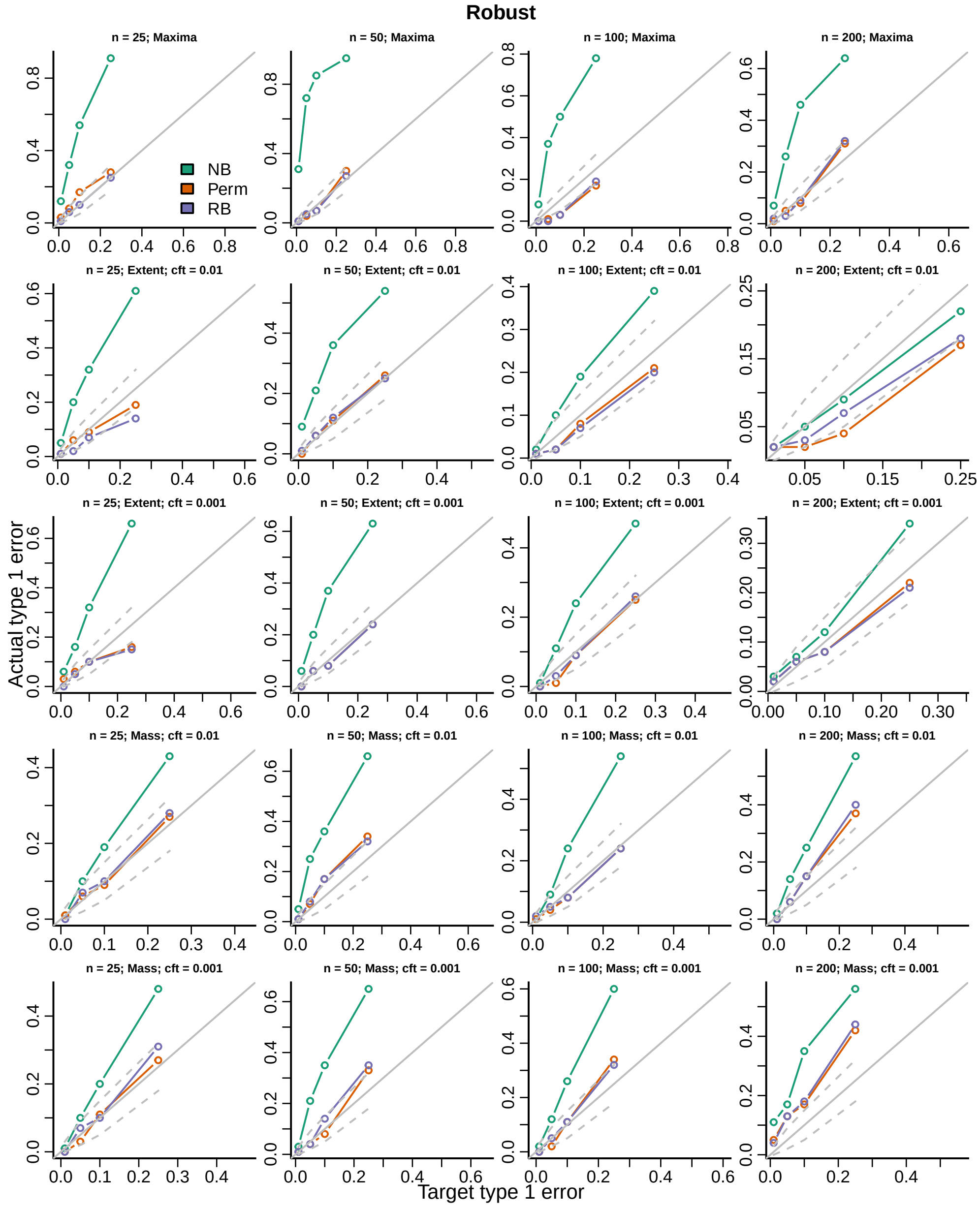


Figure S10: Actual versus target type 1 error rates for the three inference procedures considered for testing the marginal distribution of each topological feature (TF) of the robust test statistics image.

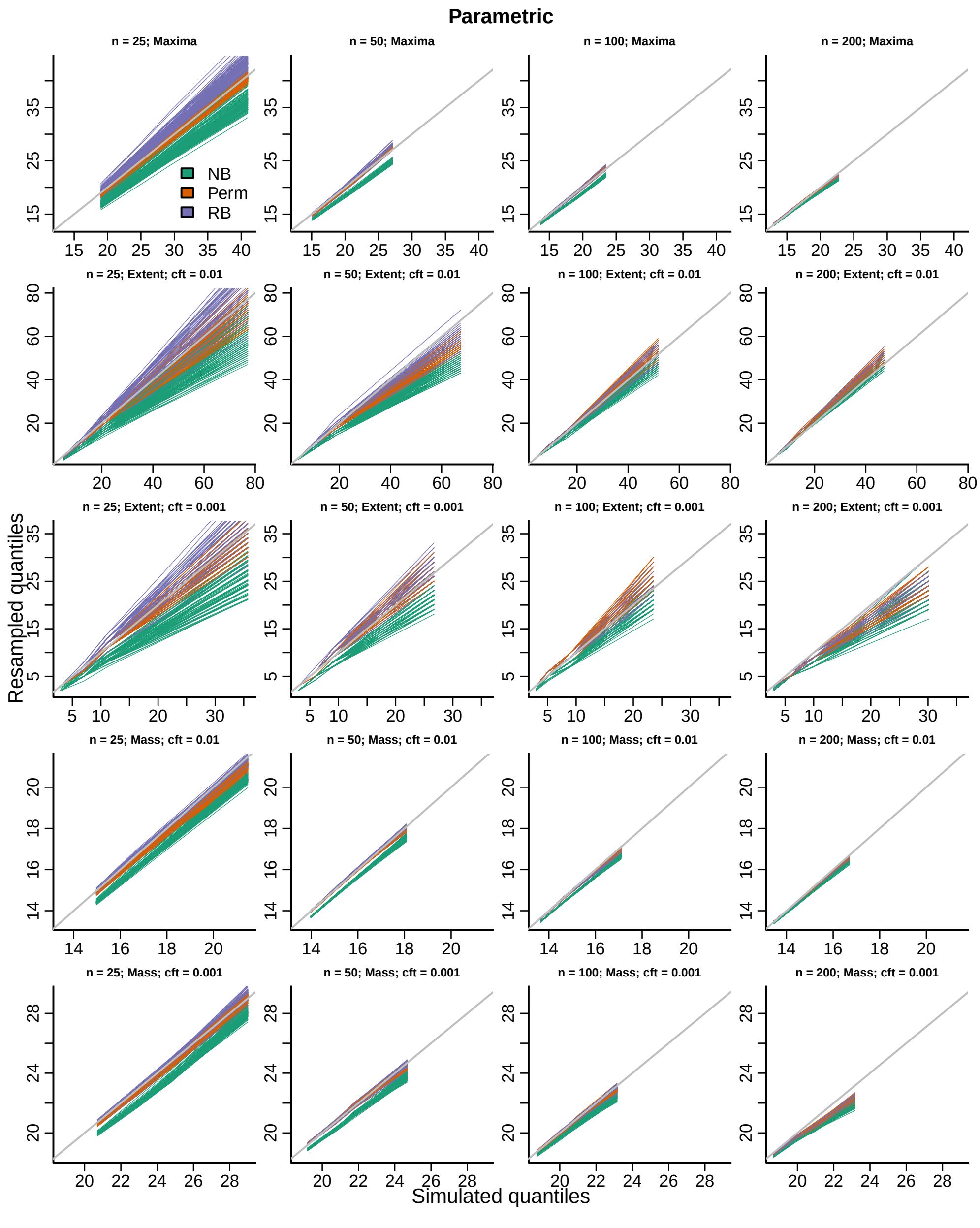


Figure S11: QQ-plots for the three inference procedures considered for the marginal distribution of each topological feature (TF) of the parametric test statistics image.

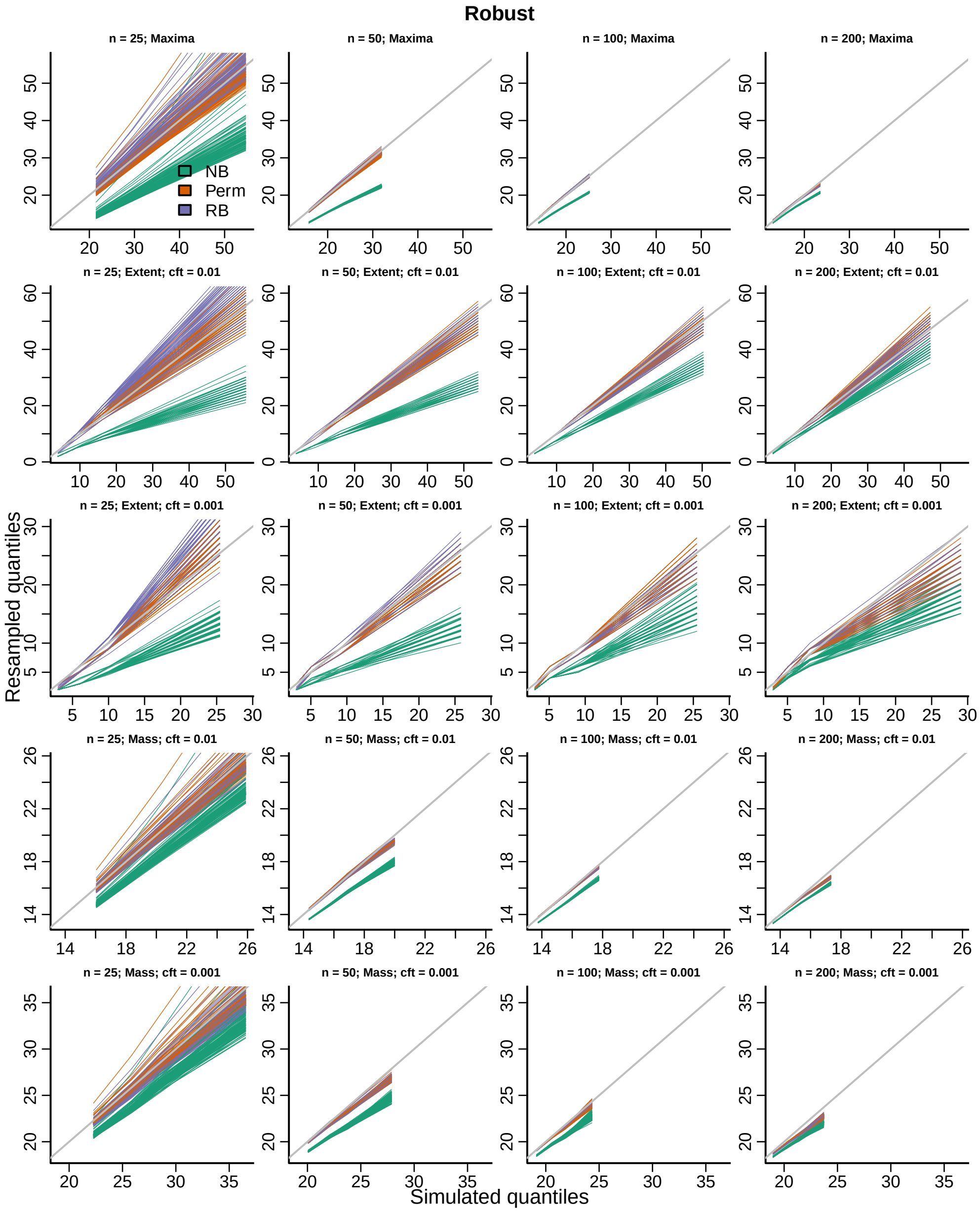


Figure S12: QQ-plots for the three inference procedures considered for the marginal distribution of each topological feature (TF) of the robust test statistics image.

##### S3.2.2 Global distributions

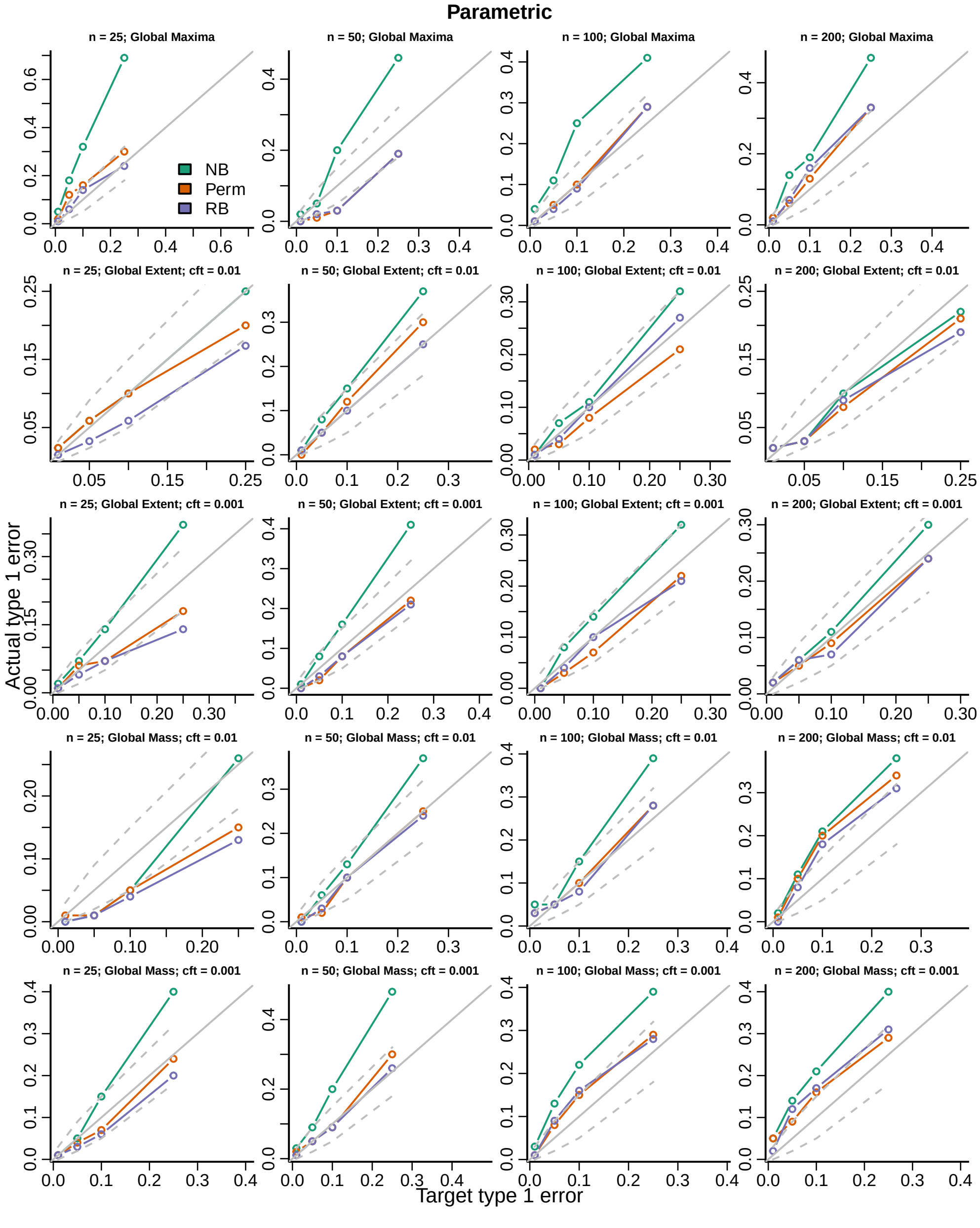


Figure S13: Actual versus target type 1 error rates for the three inference procedures considered for testing the distribution of the global maximum of each topological feature (TF) of the parametric test statistics image.

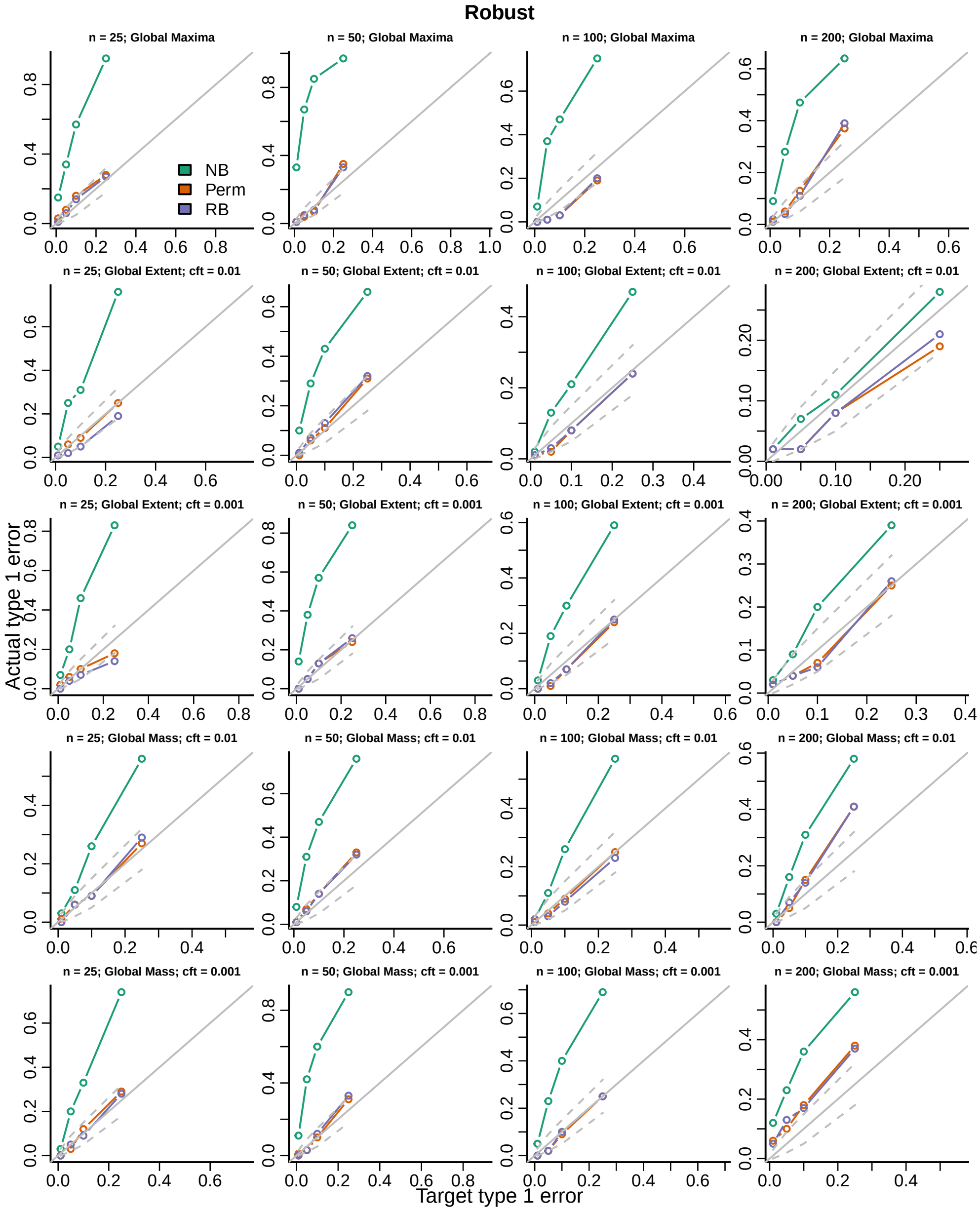


Figure S14: Actual versus target type 1 error rates for the three inference procedures considered for testing the distribution of the global maximum of each topological feature (TF) of the robust test statistics image.

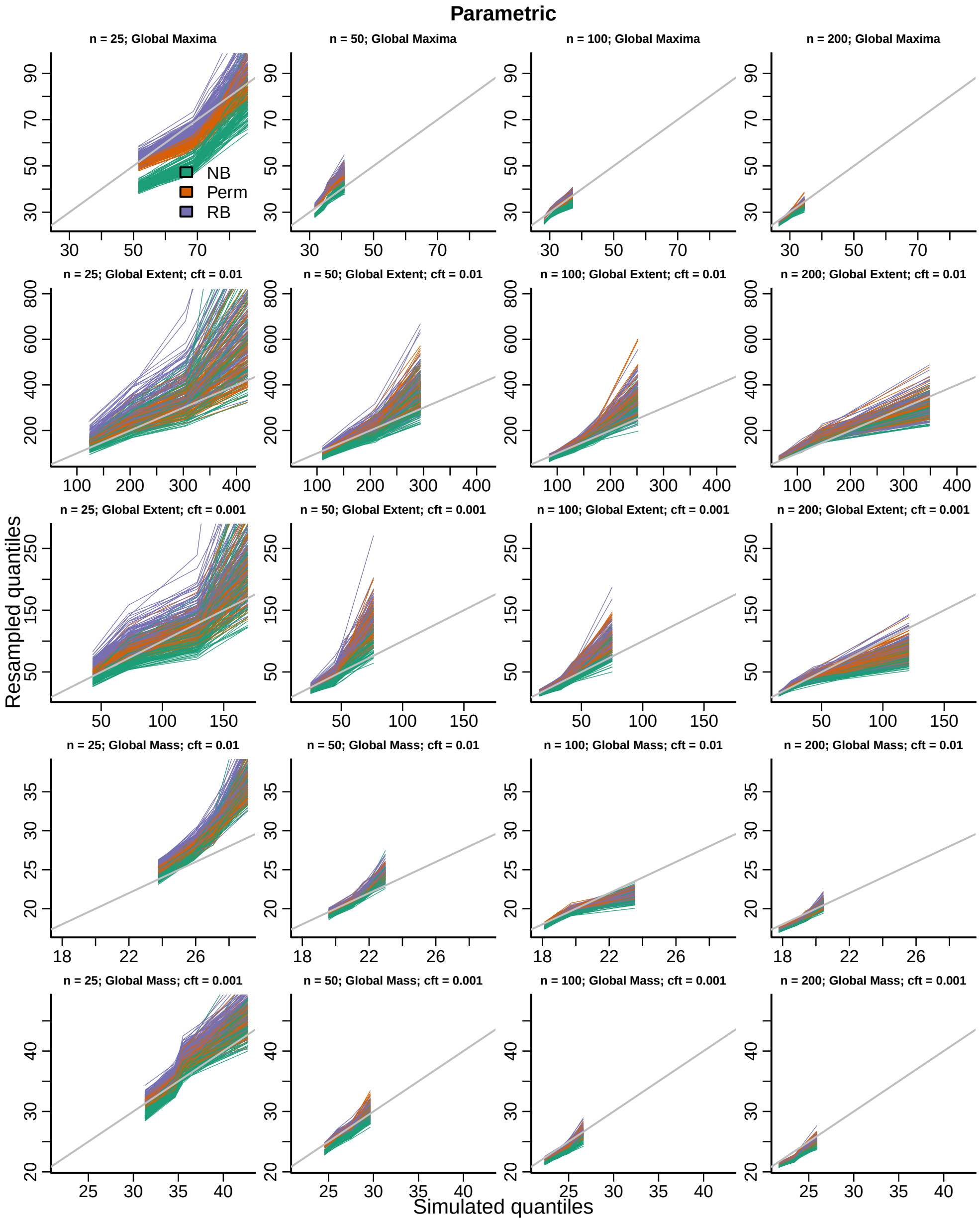


Figure S15: QQ-plots for the three inference procedures considered for the distribution of the global maximum of each topological feature (TF) of the parametric test statistics image.

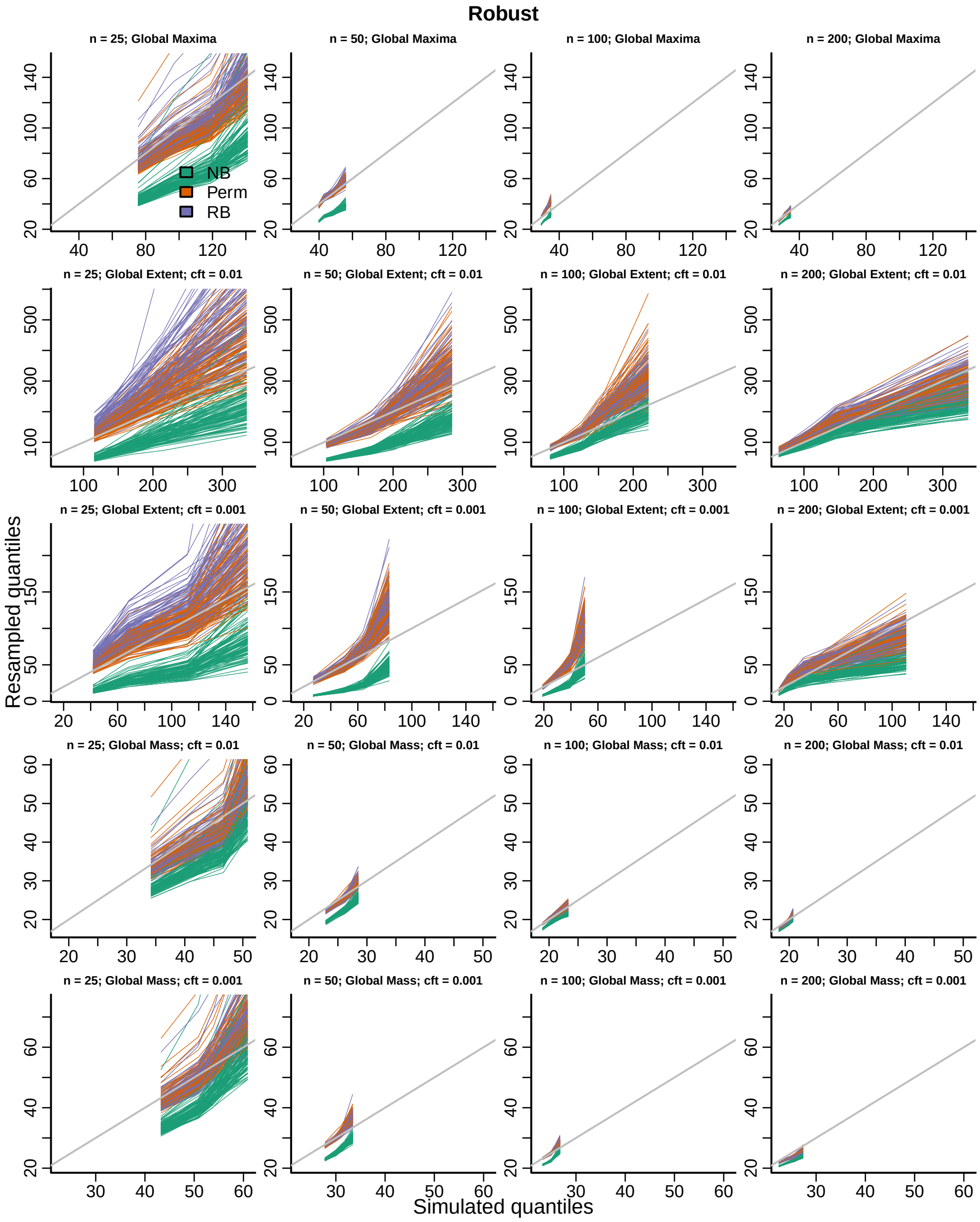


Figure S16: QQ-plots for the three inference procedures considered for the distribution of the global maximum of each topological feature (TF) of the robust test statistics image.

#### S3.3 Age continuous covariate fit with splines

The imaging data were residualized to age and other covariates using equation (13) in the main paper, then bootstrap samples of the residuals were modeled with age using splines on 4 degrees of freedom. the test was for the nonlinear effect of age over the linear age effect on 3 degrees of freedom.

##### S3.3.1 Marginal distribution

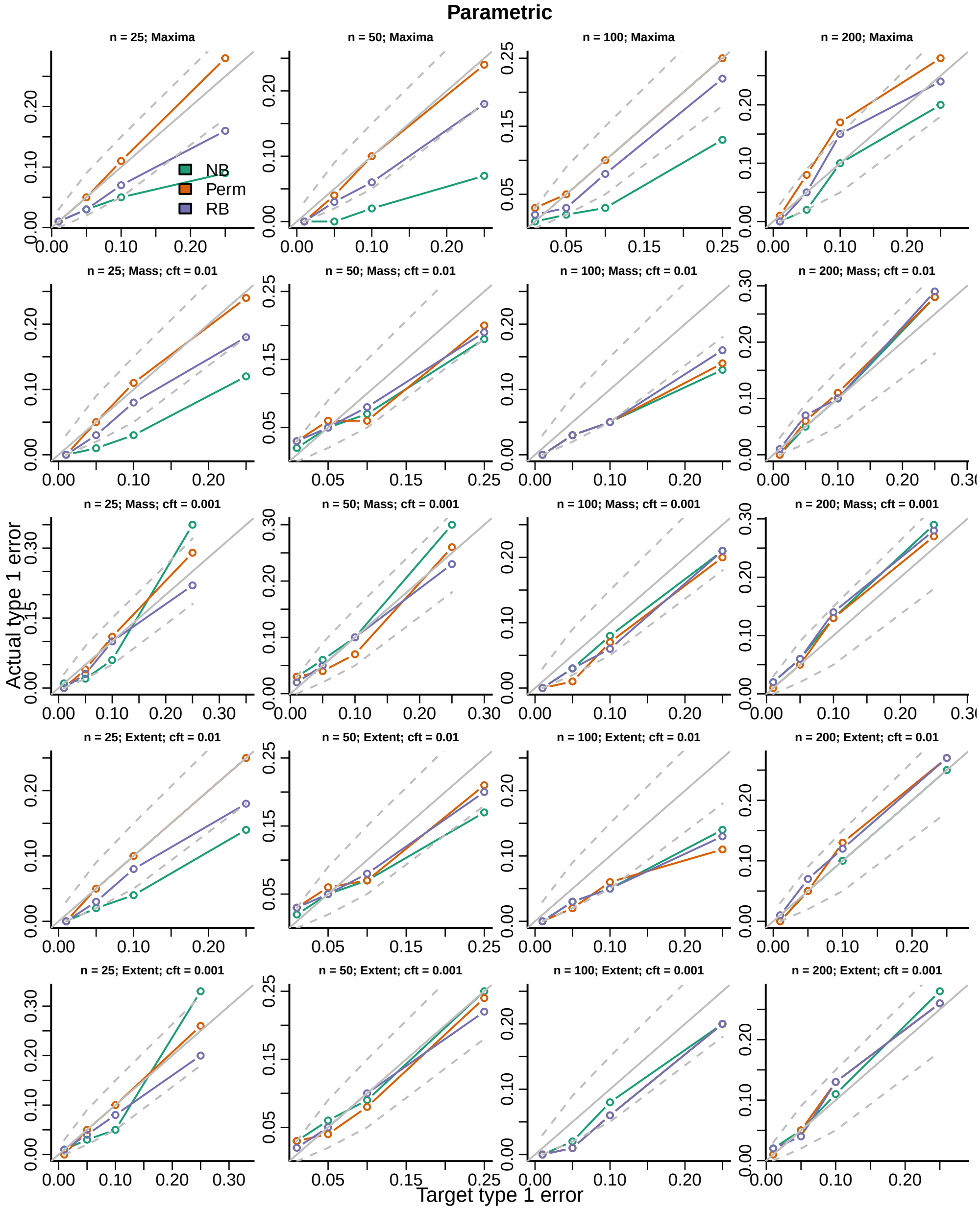


Figure S17: Actual versus target type 1 error rates for the three inference procedures considered for testing the marginal distribution of each topological feature (TF) of the parametric test statistics image.

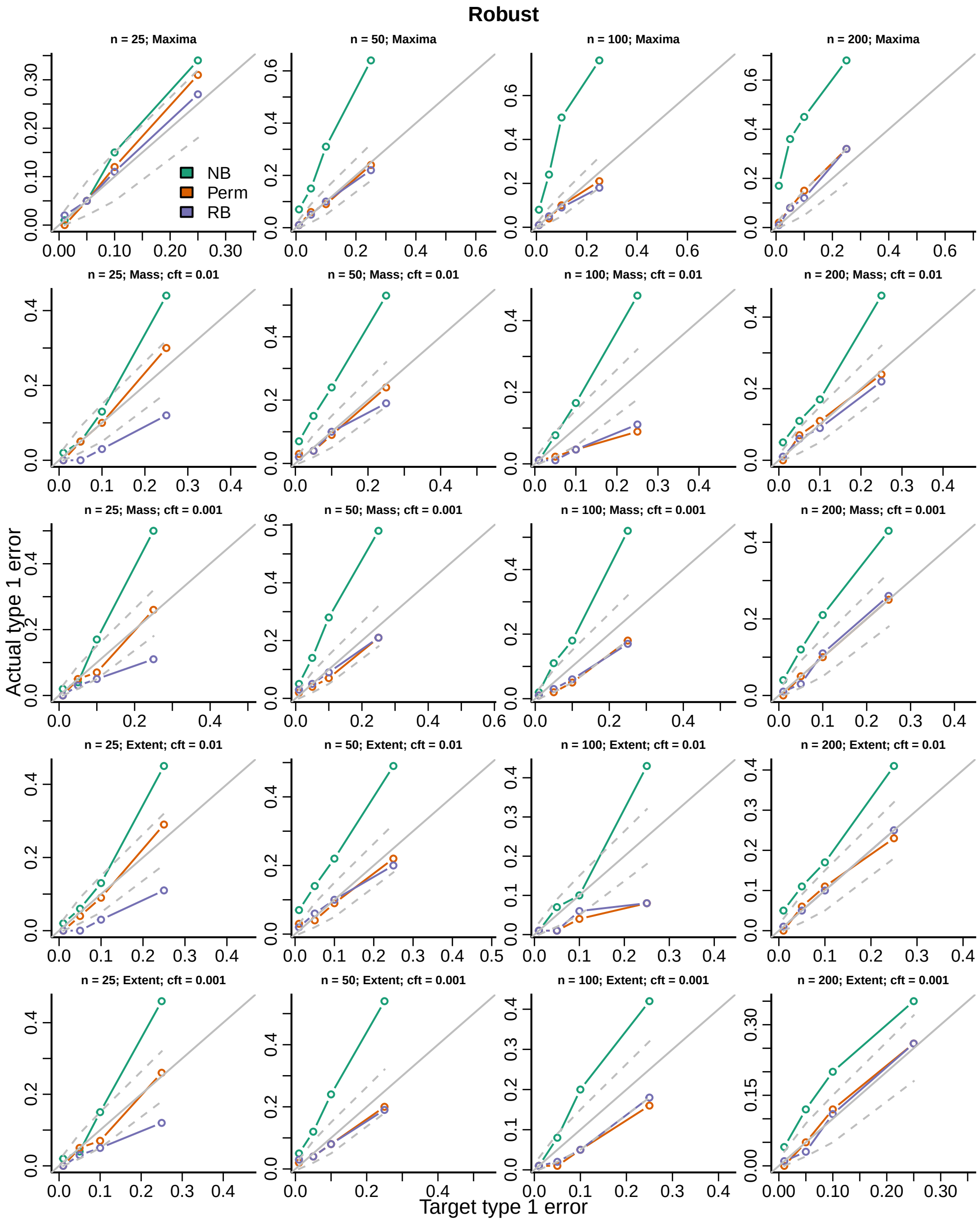


Figure S18: Actual versus target type 1 error rates for the three inference procedures considered for testing the marginal distribution of each topological feature (TF) of the robust test statistics image.

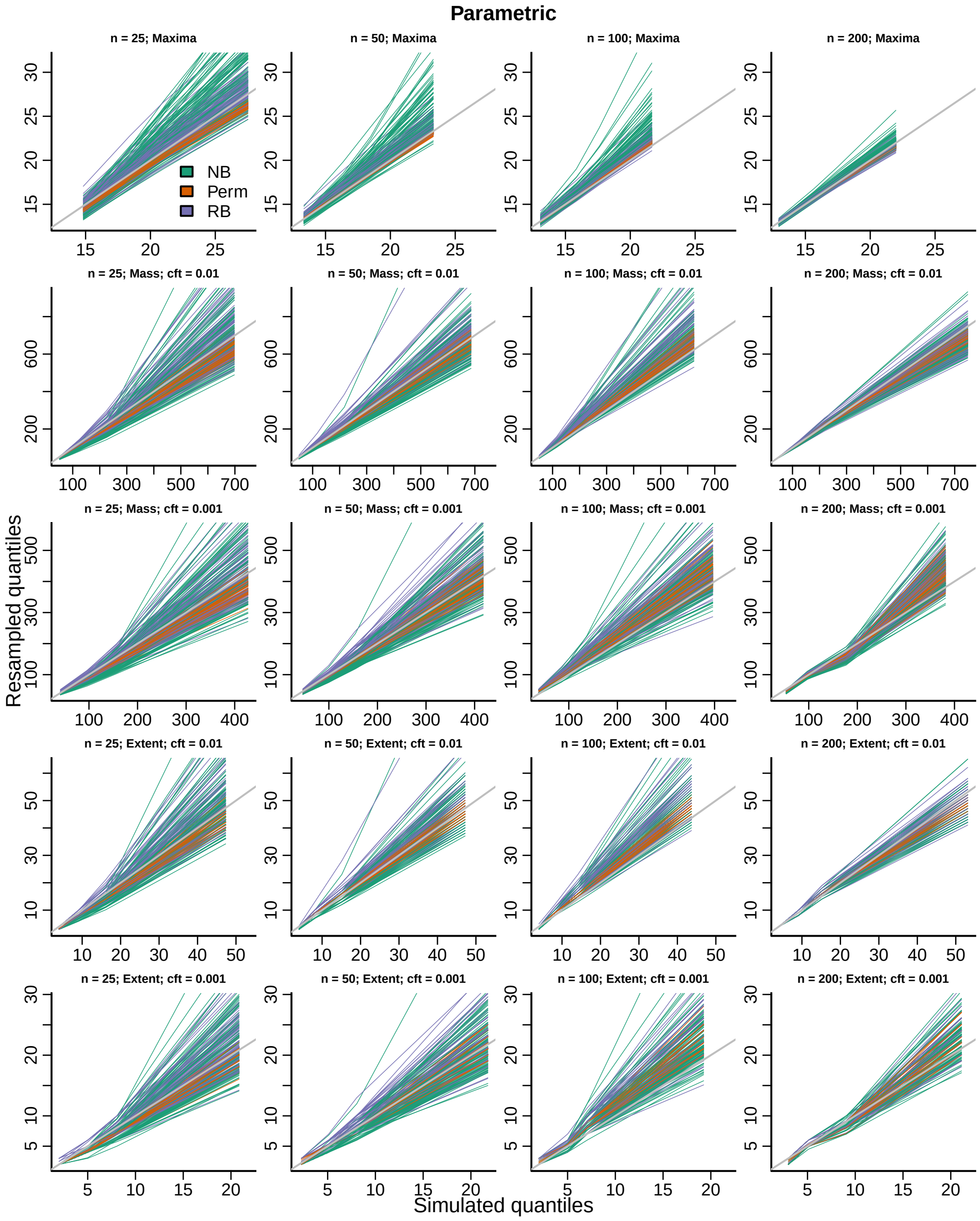


Figure S19: QQ-plots for the three inference procedures considered for the marginal distribution of each topological feature (TF) of the parametric test statistics image.

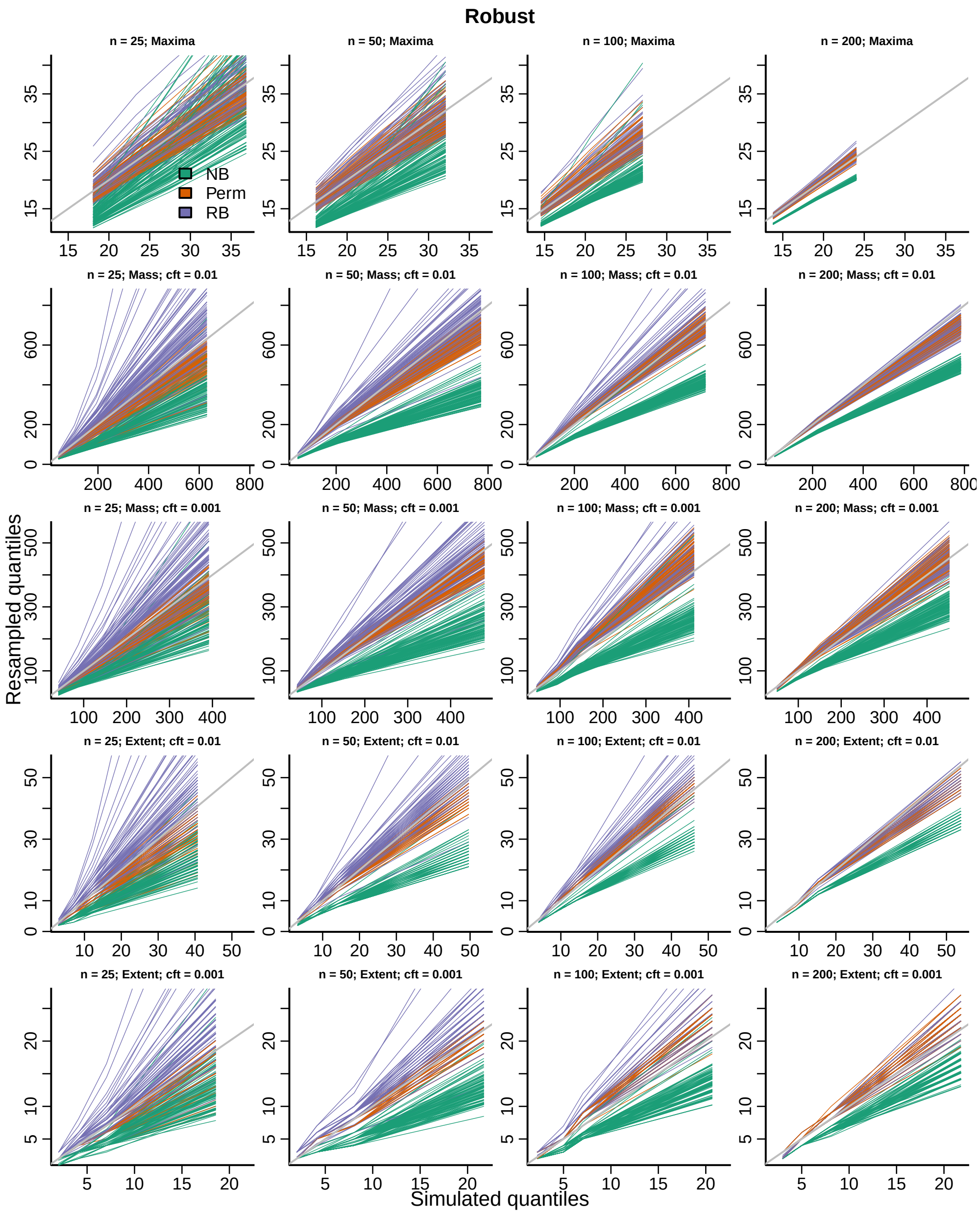


Figure S20: QQ-plots for the three inference procedures considered for the marginal distribution of each topological feature (TF) of the robust test statistics image.

##### S3.3.2 Global distributions

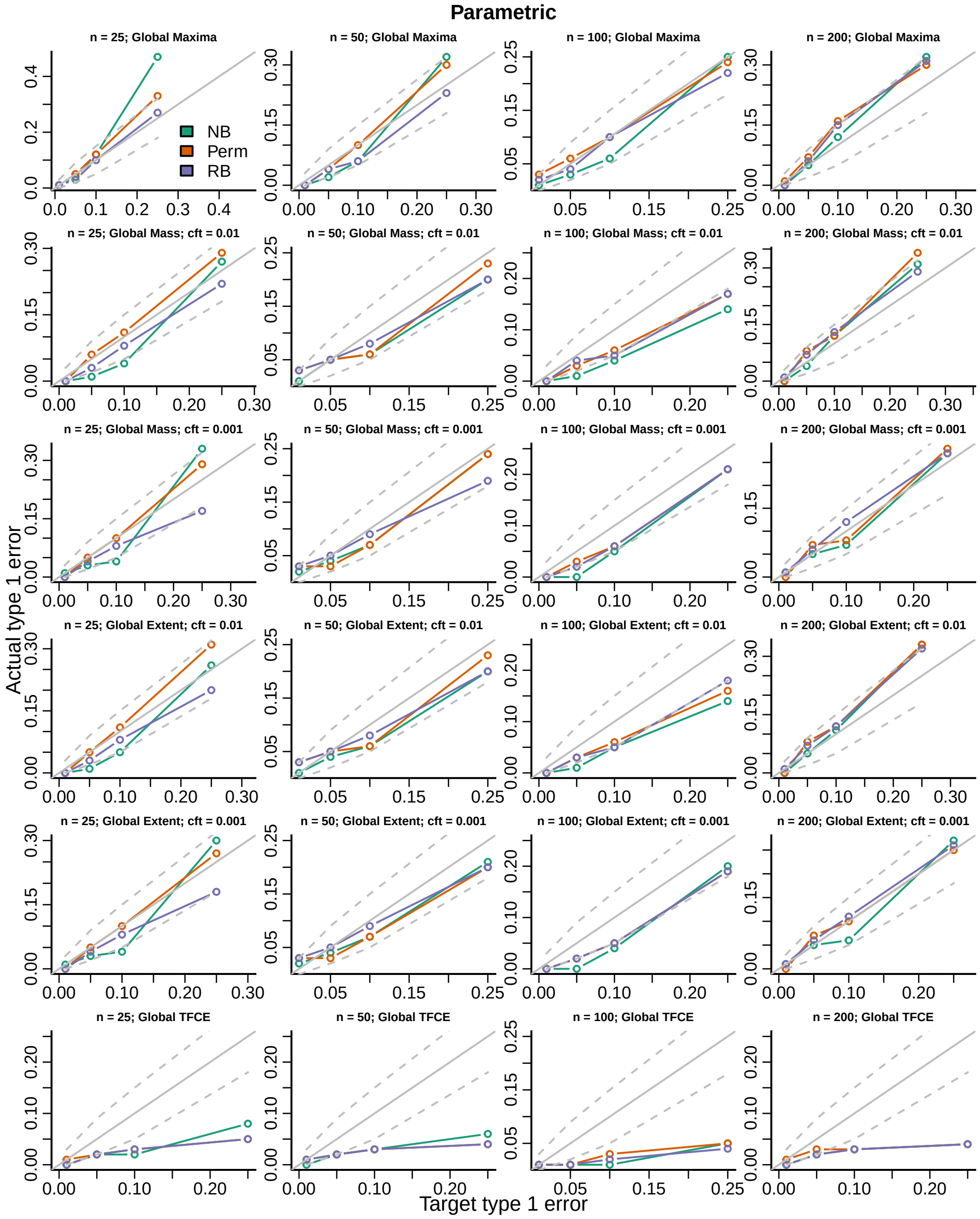


Figure S21: Actual versus target type 1 error rates for the three inference procedures considered for testing the distribution of the global maximum of each topological feature (TF) of the parametric test statistics image.

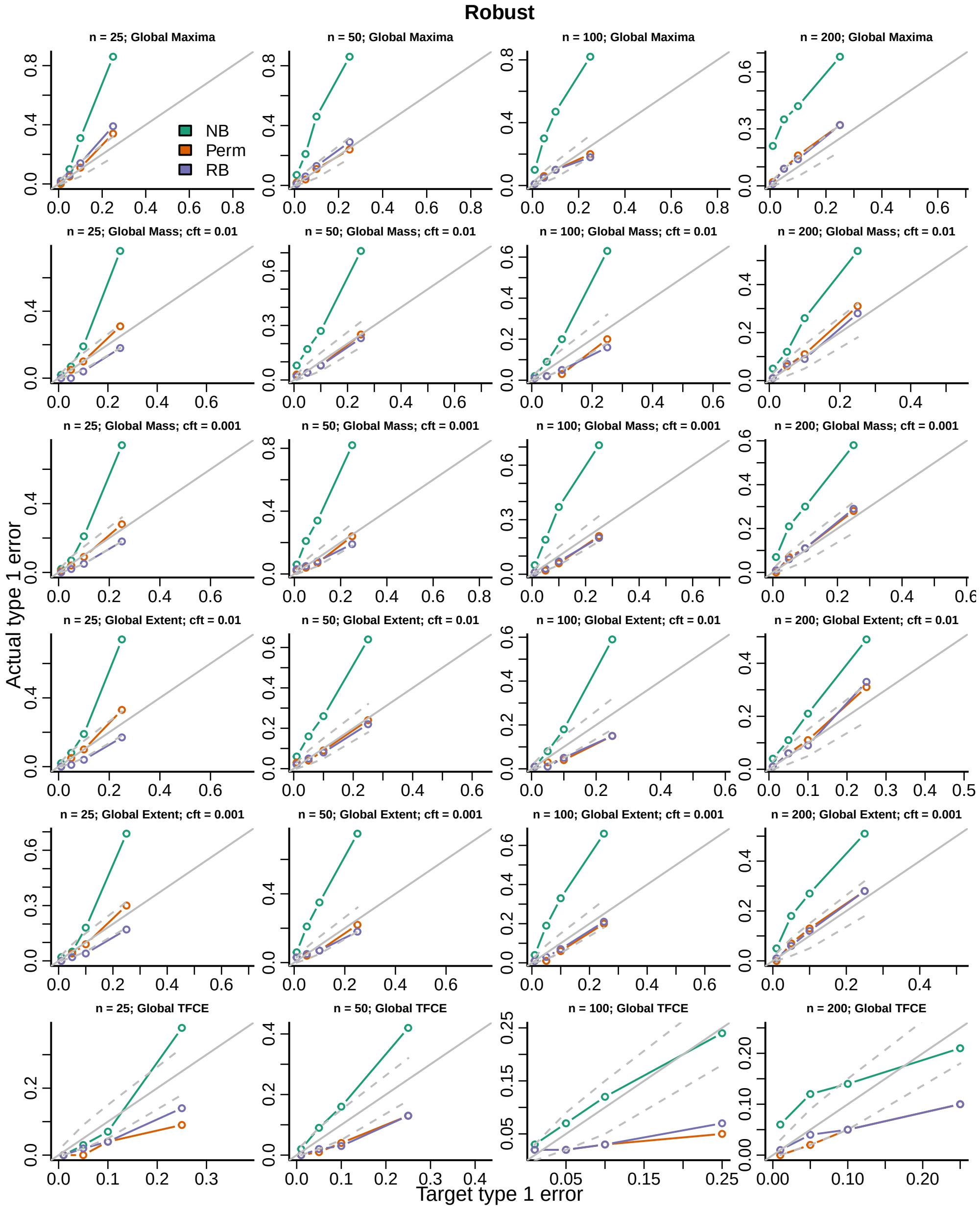


Figure S22: Actual versus target type 1 error rates for the three inference procedures considered for testing the distribution of the global maximum of each topological feature (TF) of the robust test statistics image.

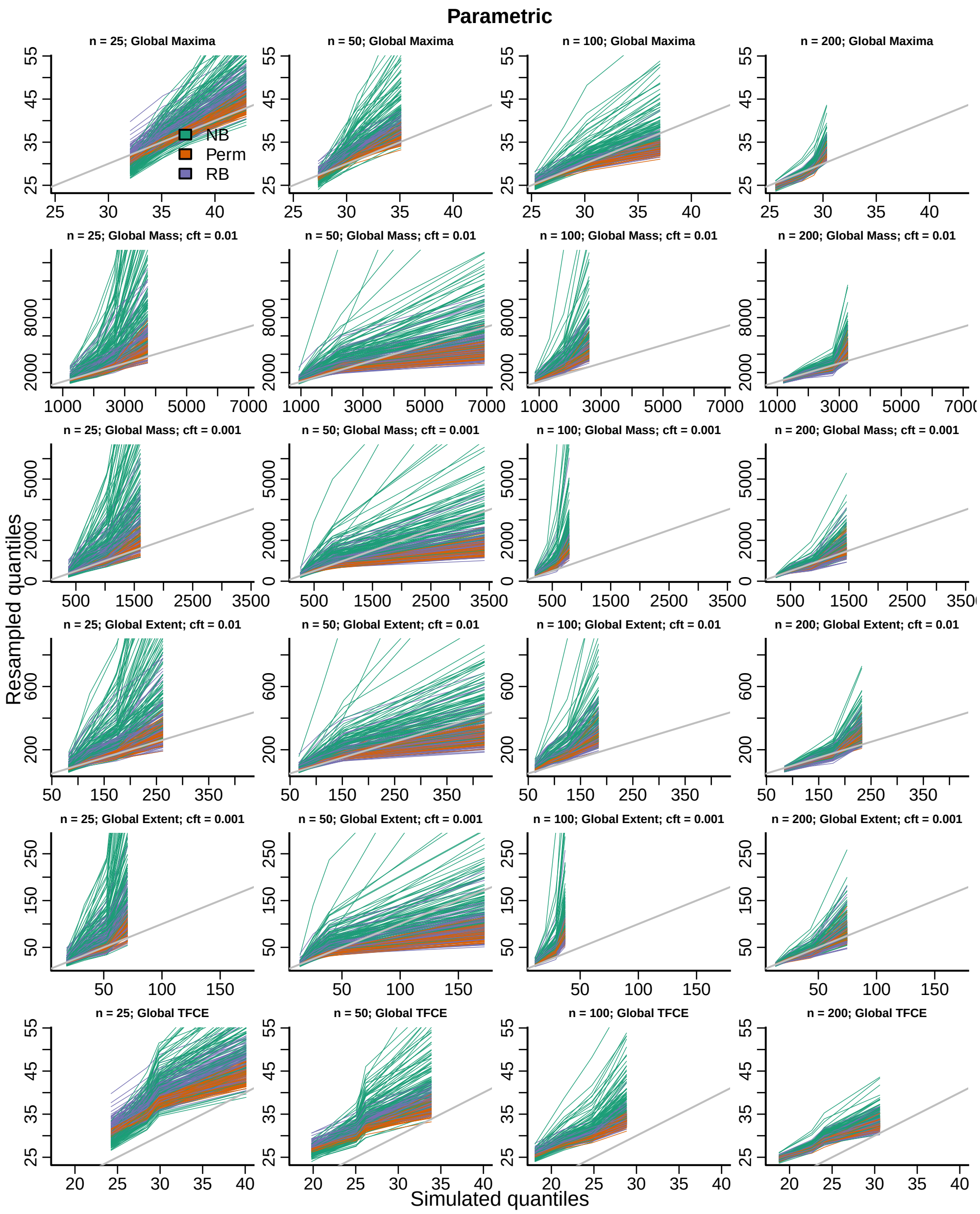


Figure S23: QQ-plots for the three inference procedures considered for the distribution of the global maximum of each topological feature (TF) of the parametric test statistics image.

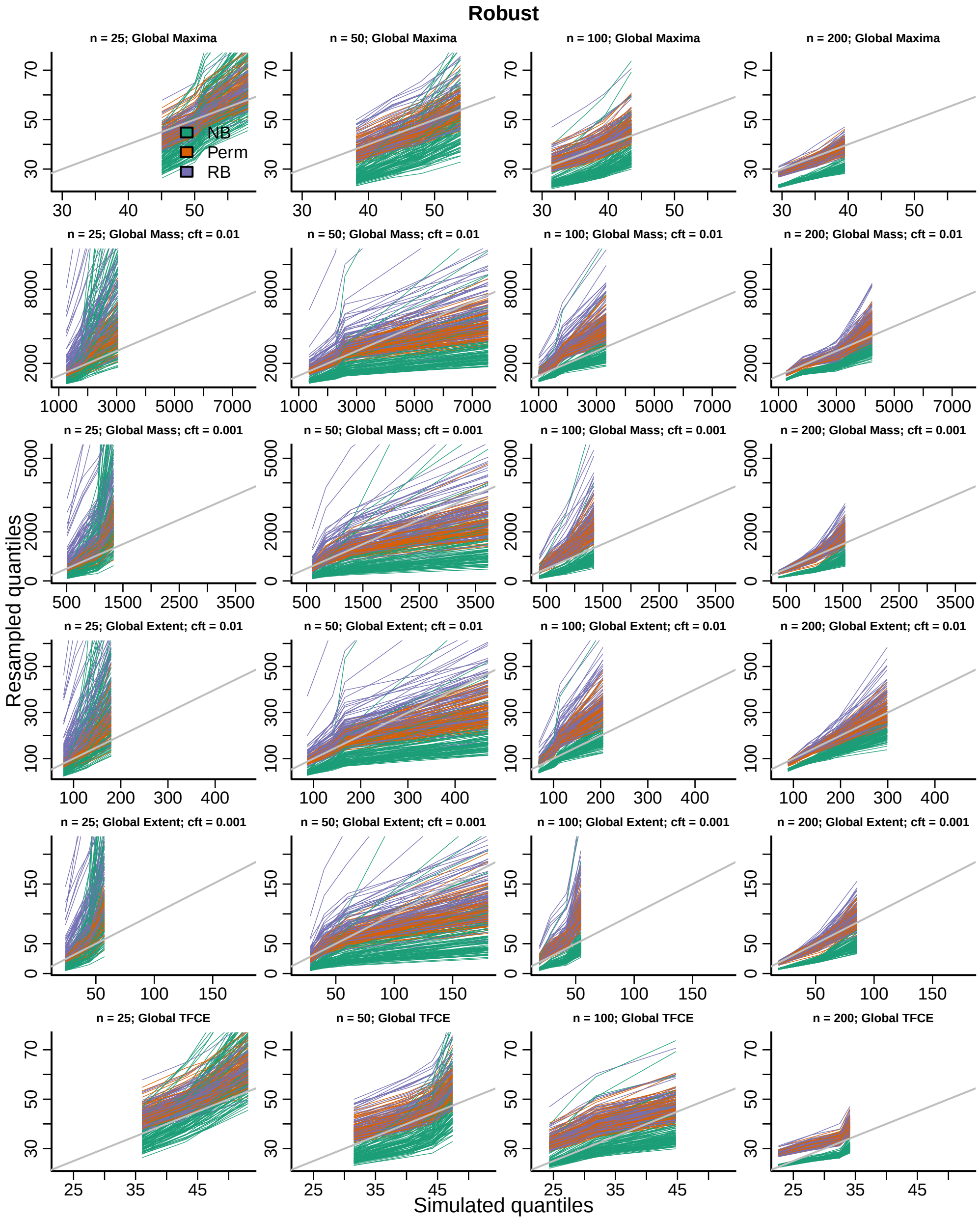


Figure S24: QQ-plots for the three inference procedures considered for the distribution of the global maximum of each topological feature (TF) of the robust test statistics image.
